# Crystal structure and RNA-binding properties of an Hfq homolog from the deep-branching Aquificae: Conservation of the lateral RNA-binding mode

**DOI:** 10.1101/078733

**Authors:** Kimberly A Stanek, Jennifer P West, Peter S Randolph, Cameron Mura

**Affiliations:** Department of Chemistry, University of Virginia, 409 McCormick Road, Charlottesville, VA, 22904, USA

**Keywords:** Hfq, Sm protein, RNA, *Aquifex aeolicus*, hexamer, evolution

## Abstract

**Synopsis:** The structure of an Hfq homolog from the deep-branching thermophilic bacterium *Aquifex aeolicus*, determined to 1.5-Å resolution both in *apo* form and bound to a uridine-rich RNA, reveals a conserved, pre-organized RNA-binding pocket on the lateral rim of the Hfq hexamer.

**Abstract:** The host factor Hfq, as the bacterial branch of the Sm family, is an RNA-binding protein involved in post-transcriptional regulation of mRNA expression and turnover. Hfq facilitates pairing between small regulatory RNAs (sRNA) and their corresponding mRNA targets by binding both RNAs and bringing them into close proximity. Hfq homologs self-assemble into homo-hexameric rings, with at least two distinct surfaces that bind RNA. Recently, another binding site—dubbed the ‘lateral rim’—has been implicated in sRNA•mRNA annealing; the RNA-binding properties of this site appear to be rather subtle, and its degree of evolutionary conservation is unknown. An Hfq homolog has been identified in the phylogenetically deep-branching thermophile *Aquifex aeolicus* (*Aae*), but little is known about the structures and functions of Hfq from basal bacterial lineages such as the Aquificae. Thus, we have cloned, overexpressed, purified, crystallized, and biochemically characterized *Aae* Hfq. We have determined the structures of *Aae* Hfq in space-groups *P*1 and *P*6, both to 1.5 Å resolution, and we have discovered nanomolar-scale binding affinities for uridine- and adenosine-rich RNAs. Co-crystallization with U_6_ RNA reveals that the outer rim of the *Aae* Hfq hexamer features a well-defined binding pocket that is selective for uracil. This *Aae* Hfq structure, combined with biochemical and biophysical characterization of the homolog, reveals deep evolutionary conservation of the lateral RNA-binding mode, and lays a foundation for further studies of Hfq-associated RNA biology in ancient bacterial phyla.

## 1. Introduction

The bacterial protein Hfq, initially identified as an *E. coli* host factor required for the replication of RNA bacteriophage Qβ (Franze de Fernandez *et al.*, 1968, Franze de Fernandez *et al.*, 1972), is now known to play a central role in the post-transcriptional regulation of gene expression and mRNA metabolism (Vogel & Luisi, 2011, Sauer, 2013, Updegrove *et al.*, 2016). Hfq has been linked to many RNA-regulated cellular pathways, including stress response (Sledjeski *et al.*, 2001, Zhang *et al.*, 2002, Fantappie *et al.*, 2009), quorum sensing (Lenz *et al.*, 2004), and biofilm formation (Mandin & Gottesman, 2010, Mika & Hengge, 2013). The diverse cellular functions of Hfq stem from its fairly generic role in binding small, non-coding RNAs (sRNA) and facilitating base-pairing interactions between these regulatory sRNAs and target mRNAs. A given sRNA might either upregulate (Soper *et al.*, 2010) or downregulate (Ikeda *et al.*, 2011) one or more target mRNAs via distinct mechanisms. For example, the sRNA RhyB downregulates several Fur-responsive genes under iron-limiting conditions (Masse & Gottesman, 2002), whereas the DsrA, RprA and ArcZ sRNAs stimulate translation of *rpoS* mRNA, encoding the stationary-phase σ^s^ factor (Soper *et al.*, 2010). In general, Hfq is required for cognate sRNA•mRNA pairings to be productive, and abolishing Hfq function typically yields pleiotropic phenotypes, including diminished viability (Fantappie *et al.*, 2009, Vogel & Luisi, 2011).

Hfq is the bacterial branch of the Sm superfamily of RNA-associated proteins (Mura *et al.*, 2013). Eukaryotic Sm and Sm-like (LSm) proteins act in intron splicing and other mRNA-related processing pathways (Will & Luhrmann, 2011, Tharun, 2009, Tycowski *et al.*, 2006), while the cellular functions of Sm homologs in the archaea remain unclear. Though the biological functions and amino acid sequences of Sm proteins vary greatly, the overall Sm fold is conserved across all three domains of life: five antiparallel β-strands form a highly bent β-sheet, often preceded by an *N*-terminal α-helix (Fig 1; (Kambach *et al.*, 1999)). Sm proteins typically form cyclic oligomers via hydrogen bonding between the β4⋯β5′ (edge) strands of monomers in a head–tail manner, yielding a toroidal assembly of six (Hfq) or seven (other Sm) subunits (Mura *et al.*, 2013). Hfq (and other Sm) rings can further associate into head–head and head–tail stacked rings, as well as polymeric assemblies (Arluison *et al.*, 2006); any *in vivo* relevance of double-rings and other higher-order species remains unclear (Mura *et al.*, 2013). The oligomerization mechanism also varies: Sm-like archaeal proteins (SmAPs) and Hfq homologs spontaneously self-assemble into stable homo-heptameric or homo-hexameric rings (respectively) that resist chemical and thermal denaturation, whereas eukaryotic Sm hetero-heptamers form via a chaperoned biogenesis pathway. This intricate assembly pathway (Fischer *et al.*, 2011) involves staged interactions with single–stranded RNA (e.g. small nuclear RNAs of the spliceosomal snRNPs), such that RNA threads through the central pore of the Sm ring (Leung *et al.*, 2011). In contrast, Hfq hexamers expose two distinct RNA-binding surfaces (Mikulecky *et al.*, 2004), termed the ‘proximal’ and ‘distal’ (with respect to the α-helix) faces of the ring. These two surfaces can bind RNA independently and simultaneously (Wang *et al.*, 2013), with different RNA sequence specificities along each face.

**Figure 1.**
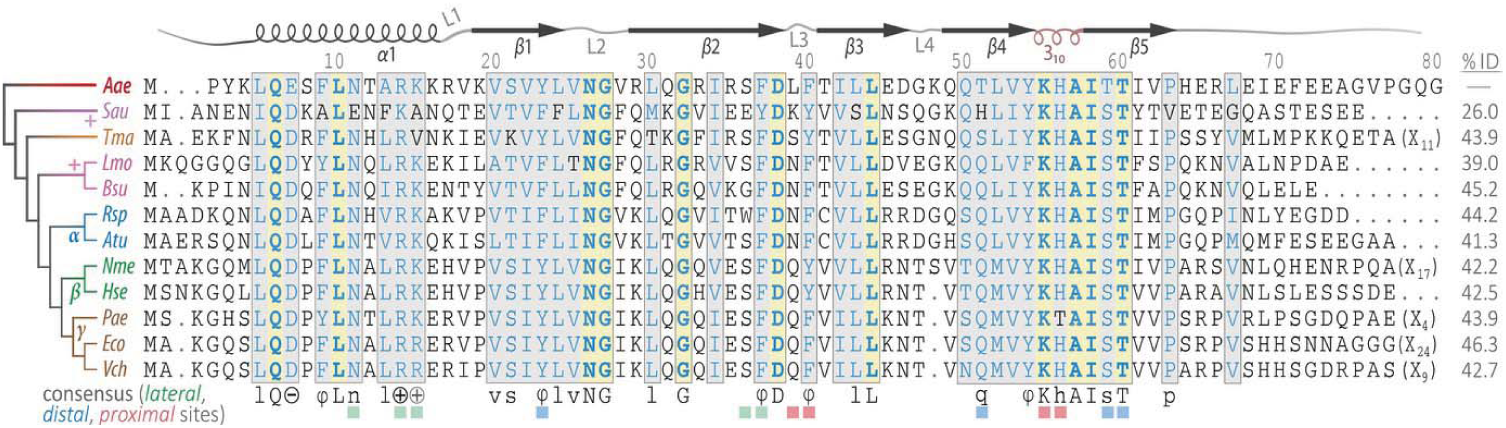
Multiple sequence alignment of *Aae* Hfq and some representative homologues. Sequence analysis of several Hfq homologues, characterized from various phyla, reveals conservation of key amino acids comprising Hfq’s three distinct RNA-binding regions (*distal*, *proximal*, *lateral*). The *Aae* Hfq sequence is numbered at the top, and secondary structural elements are drawn based on the *Aae* crystal structures reported herein; helices are schematized as spirals, strands as arrows, and numbered loop labels are shown (a short 3_10_ helix forms loop L5, coloured brown). Strictly identical amino acids are in bold blue text on a yellow background, while sites with highly similar residues are highlighted with a grey background; these blocks of partially conserved residues are also lightly boxed. In the consensus sequence shown at the bottom, uppercase letters indicate strict identity and lowercase letters correspond to physicochemically equivalent residues that meet the similarity threshold (≥85% sites in a given column). Residues known to contact RNA at the proximal, distal, or lateral sites are marked with red, blue, or green square symbols, respectively. Note the high level of conservation of residues involved in all three RNA-binding sites. In addition to *Aae* Hfq (of the phylum *Aquificae*), the twelve aligned sequences include (*i*) three Hfq homologs from the mostly Gram–positive Firmicutes (*Sau*, *Lmo*, *Bsu*), (*ii*) a homologue from the ancient phylum *Thermotogae* and (*iii*) several characterized Hfq orthologues from the α–, β– and γ–proteobacteria. The relationships between these species are indicated in the dendrogram (left), obtained during the progressive alignment calculation and coloured so as to highlight phylum-level differences. The genus*/*species and sequence accession codes [GenBank] follow: *Aae*, *A. aeolicus* [AAC06479.1]; *Sau*, *Staphylococcus aureus* [ADC37472.1]; *Tma*, *T. maritima* [AGL49448.1]; *Lmo*, *Listeria monocytogenes* [CBY70202.1]; *Bsu*, *Bacillus subtilis* [BAM57957.1]; *Rsp*, *Rhodobacter sphaeroides* [A3PJP5.1]; *Atu*, *Agrobacterium tumefaciens* [EHH08904.1]; *Nme*, *Neisseria meningitidis* [P64344.1]; *Hse*, *Herbaspirillum seropedicae* [ADJ64436.1]; *Pae*, *P. aeruginosa* [B3EWP0.1]; *Eco*, *E. coli* [BAE78173.1]; and *Vch*, *Vibrio cholerae* [A5F3L7.1].

The proximal face of Hfq preferentially binds uridine-rich single-stranded RNA (ssRNA) in a manner that is well-conserved amongst Gram-positive (Schumacher *et al.*, 2002, Kovach *et al.*, 2014) and Gram-negative bacteria (Weichenrieder, 2014). The binding region, located near the pore, consists of six equivalent ribonucleotide-binding pockets, and can thus accommodate a six-nucleotide segment of ssRNA. Each uracil base π-stacks between a conserved aromatic side-chain (Phe or Tyr) from the L3 loops of adjacent monomers (e.g., F42 in *E. coli*; Fig 1), and nucleobase specificity is achieved via hydrogen bonding between Q8 and the exocyclic O2 of each uracil (unless otherwise noted, residue numbers refer to the *E. coli* Hfq sequence). A key physiological function of the proximal face of Hfq is thought to be the selective binding of the U–rich 3′-termini of sRNAs, resulting from *ρ*–independent transcription termination (Wilson & von Hippel, 1995). Hfq’s recognition of these 3′ ends is facilitated by H57 of the L5 loop (‘3_10_ helix’ in Fig 1), which is well-positioned to interact with the unconstrained, terminal 3′-hydroxyl group (Sauer & Weichenrieder, 2011, Schulz & Barabas, 2014). This mode of recognition may also explain the ability of Hfq to bind specifically to sRNAs over DNA, or other RNAs.

In contrast to the uracil-binding proximal region, the distal face of Hfq preferentially binds adenine-rich RNA, with the mode of binding varying between Gram-negative and Gram-positive species. Hfq homologs from Gram-negative bacteria specifically recognize RNAs with a tri-nucleotide motif, denoted (A–R–N)_*n*_, where A=adenine, R=purine, N=any nucleotide; this recognition element was recently refined to be a more restrictive (A–A–N)_*n*_ motif (Robinson *et al.*, 2014). AAN–containing RNAs bind to a large surface region on the distal face, which can accommodate up to 18 nucleotides of a ssRNA (Link *et al.*, 2009), and such RNAs are recognized in a tripartite manner: (*i*) the first *A site* is formed by residues between the β2 and β4 strands of one monomer (E33 ensures adenine specificity); (*ii*) the second *A site* lies between the β2 strands of adjacent subunits, and includes a conserved Y25 (Fig 1) that engages in π-stacking interactions; and (*iii*) a nonspecific nucleotide (N)–binding site bridges to the next A–A pocket. In contrast to this recognition mechanism, the distal face of Gram-positive Hfq recognizes a bipartite adenine–linker (AL)_*n*_ motif. This structural motif features an A site that is similar to the first A-site of Gram-negative bacteria; in addition, a nonspecific nucleotide-binding pocket acts as a linker (*L*) site, allowing 12 nucleotides to bind in a circular fashion atop this face of the hexamer (Horstmann *et al.*, 2012, Someya *et al.*, 2012). The ability of the distal face to specifically bind A-rich regions, such as the long, polyadenylated 3′-tails of mRNAs (Folichon *et al.*, 2003), leads to several links between Hfq and mRNA degradation*/*turnover pathways (Mohanty *et al.*, 2004, Bandyra & Luisi, 2013, Regnier & Hajnsdorf, 2013). The general capacity of Hfq to independently bind RNAs at the proximal and distal sites brings these distinct RNA species into close proximity, as part of an sRNA•Hfq•mRNA ternary complex. Indeed, a chief cellular role of Hfq is the productive annealing of RNA strands in this manner, for whatever downstream physiological purpose (be it stimulatory or inhibitory).

Independent binding of RNAs at the proximal/distal sites elucidates only part of what is known about Hfq’s RNA-related activities. For instance, Hfq has been shown to protect internal regions of sRNA (Balbontín *et al.*, 2010, Ishikawa *et al.*, 2012, Updegrove & Wartell, 2011, Zhang *et al.*, 2002) and to reduce the thermodynamic stability 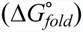 of some RNA hairpins (Robinson *et al.*, 2014), but current mechanistic models for Hfq activity do not account for all of these properties. In addition, recent studies have identified a new RNA-binding site on the Hfq ring, beyond the proximal and distal sites (Sauer, 2013). This third site, located on the outer rim of the Hfq toroid and presaged in RNA-binding studies a decade ago (Sun & Wartell, 2006), is variously termed the ‘lateral’, ‘rim’ or ‘lateral rim’ site (the terms are used synonymously herein). Mutational analyses reveal that an arginine-rich patch near the *N*-terminal α-helix, containing the segment R^16^R^17^E^18^R^19^ in *E. coli*, facilitates rapid annealing of Hfq-bound mRNAs and sRNAs (Panja *et al.*, 2013). These arginine residues, along with conserved aromatic (F/Y39; ‘φ’ in Fig 1) and basic (K47) residues, look to be vital for the binding of full-length sRNAs to Hfq (Sauer *et al.*, 2012). Further understanding of the precise mechanism of RNA binding to the lateral site (and any base specificity at that site) has been hindered by a lack of structural information on Hfq_*rim*_⋯RNA interactions. A recent crystal structure of *E. coli* Hfq complexed with the full-length riboregulatory sRNA RydC (a regulator of biofilms and some mRNAs) revealed a potential binding pocket formed by N13, R16, R17 and F39, and capable of accommodating two nucleotides of uridine (Dimastrogiovanni *et al.*, 2014); however, the exact positioning and geometry of the nucleotides were not discernible at the resolution (3.5 Å) of that model.

Our current mechanistic knowledge of Hfq⋯RNA interactions is based on homologs from proteobacterial species, particularly the γ-proteobacteria *E. coli* and *Pseudomonas aeruginosa*; structural information about nucleotide-binding at the lateral site is available only from these two species. We do not know if the rim RNA-binding mode is conserved in homologs from other bacterial species, or perhaps even more broadly (in archaeal and eukaryotic lineages). Hfq orthologs from phylogenetically deep-branching bacteria, such as *Aquifex aeolicus* (*Aae*), may help clarify the degree of conservation of Hfq’s various RNA-binding surfaces, including the lateral rim. *Aae* Hfq has been shown, via immunoprecipitation/deep-sequencing studies, to partially restore the phenotype of a *Salmonella enterica* Hfq knock-out strain, *Δhfq* (Sittka *et al.*, 2009), but nothing else is known about the RNA-binding properties of *Aae* Hfq. Precisely positioning *Aae* within the bacterial phylogeny is difficult given, for instance, that many *Aae* genes are similar to those in ε-proteobacteria (Eveleigh *et al.*, 2013). Nevertheless, 16S rRNA and genomic sequencing data firmly place *Aae*, along with other members of the Aquificales phylum, among the deepest branches in the bacterial tree—near the bacterial/archaeal divergence. Sequence similarity to proteobacterial genes has been attributed to extensive lateral gene transfer (Oshima *et al.*, 2012, Boto, 2010); importantly, extensive lateral transfer does not seem to be an issue with Hfq homologs (Sun *et al.*, 2002), and Sm proteins likely have a single, well-defined origin (Veretnik *et al.*, 2009).

Here, we report the crystal structure and RNA-binding properties of an *A. aeolicus* Hfq ortholog. *Aae* Hfq crystallized in multiple space-groups, with both hexameric and dodecameric assemblies in the lattices. These oligomeric states were further examined in solution, via chemical crosslinking assays, analytical size-exclusion chromatography, and light-scattering experiments. We found that *Aae* Hfq binds uridine– and adenosine–rich RNAs with nanomolar affinities *in vitro*, and that the inclusion of Mg^2+^ enhances binding affinities by factors of ≈2× (A-rich) or ≈10× (U-rich). Co-crystallization of *Aae* Hfq with U_6_ RNA reveals well-defined electron density (to 1.5 Å) for at least two ribonucleotides in a rim site, suggesting that this auxiliary RNA-binding site is conserved even amongst evolutionarily ancient bacteria. Finally, comparative structural analysis reveals that (*i*) the spatial pattern of Hfq⋯ RNA interatomic contacts, which effectively defines the rim site, is preserved between *Aae* and *E. coli*, and (*ii*) the residues comprising the *Aae* Hfq rim site are pre-organized for U–rich RNA binding.

## 2. Materials and Methods

### 2.1. Cloning, expression and purification of *Aae* Hfq

The *Aae hfq* gene was cloned via the polymerase incomplete primer extension (PIPE) methodology (Klock & Lesley, 2009), using an *A. aeolicus* genomic sample as a PCR template. The T7-based expression plasmid pET-28b(+) was used, yielding a recombinant protein construct bearing an *N*-terminal His6×-tag and a thrombin-cleavable linker preceding the Hfq (Supp Fig S1a, Supp Table S1); in all, the tag and linker extended the 80-aa native sequence by 20 residues. Plasmid amplification, and *in vivo* ligation of the vector and insert, were achieved via transformation of the PIPE products into chemically competent TOP10 *E. coli* cells. Recombinant *Aae* Hfq was produced by transforming the plasmid into the BL21(DE3) *E. coli* expression strain, followed by outgrowth in Luria-Bertani media at 310 K. Finally, expression of *Aae* Hfq from the T7*lac*-based promoter was induced by the addition of 1 mM isopropyl-β-D-thiogalactoside (IPTG) when the optical density, measured at 600 nm (OD_600_) reached ≈ 0.8–1.0. Cell cultures were then incubated at 310 K, with shaking (≈230rpm), for an additional four hours, pelleted at 15,000*g* for 5 minutes at 277 K, and then stored at 253 K overnight.

Cell pellets were re-suspended in a solubilisation and lysis buffer (50 mM Tris pH 7.5, 750 mM NaCl, 0.4 mM PMSF, and 0.01 mg/ml chicken egg white lysozyme (Fisher)) and incubated at 310 K for 30 min. Cells were then mechanically lysed using a microfluidizer. To clarify cell debris, the lysate was pelleted via centrifugation at 35,000*g* for 20 min at 277 K. The supernatant from this step was then incubated at 348 K for 20 min, followed by centrifugation at 35,000*g* for 20 min; this heat-cut step was performed because most Hfq homologs examined thus far have been thermostable, and because *A. aeolicus* is a hyperthermophile (optimum growth temperature, *T*_*opt*_ ≈ 368 K (Huber & Eder, 2006)). To reduce contamination by any spurious *E. coli* nucleic acids, which have been known to co-purify with other Hfqs, the clarified supernatant from the heating step was treated with high concentrations (≈ 6 M) of guanidinium hydrochloride (GndCl), followed immediately by 0.2-µm syringe filtration.

Recombinant *Aae* Hfq was then purified via immobilized metal affinity chromatography (IMAC), using a Ni^2+^–charged iminodiacetic acid–sepharose column with an NGC (BioRad) medium-pressure liquid chromatography system. After loading the clarified supernatant from the heat-cut and GndCl treatment steps, the column was treated with four column volumes of wash buffer (50 mM Tris pH 8.5, 150 mM NaCl, 6 M GndCl, 10 mM imidazole). Next, *Aae* Hfq was eluted by applying a linear gradient, from 0–100% over 10 column volumes, of elution buffer (identical to the wash buffer, but with 600 mM imidazole). Protein-containing fractions, as assessed by the absorbance at 280 nm and chromatogram elution profiles, were then combined and dialyzed into a buffer of 25 mM Tris pH 8.0, 1 M arginine in order to remove GndCl. The protein was then dialyzed into 50 mM Tris pH 8.0, 500 mM NaCl and 12.5 mM EDTA in preparation for removal of the His6×-tag. The *Aae* Hfq sample was subjected to proteolysis with thrombin, at a 1:600 Hfq:thrombin ratio (by mass), by incubating at 315 K overnight (≈16 h), followed by application to a benzamidine affinity column to remove the thrombin. To improve sample homogeneity, *Aae* Hfq was further purified over a preparative-grade gel-filtration column containing *Superdex™ 200 Increase* resin; *Aae* Hfq eluted as a single, well-defined peak. Chromatographic steps were conducted at room temperature; lengthier incubation steps, such as dialysis, were carried out at 310 or 315 K throughout the purification, as *Aae* Hfq samples were found to be relatively insoluble over a few hours at room temperature (≈ 295 K).

*Aae* Hfq sample purity was generally assayed via SDS-PAGE gels or matrix assisted laser desorption/ionization time-of-flight mass spectrometry (MALDI-TOF MS). Samples were prepared for MALDI by diluting 1:4 (v/v) with 0.01% v/v trifluoroacetic acid (TFA) and then spotting on a steel MALDI plate in a 1:1 v/v ratio with a matrix solution (15 mg/ml sinapinic acid in 50% acetonitrile, 0.05% TFA); this mixture crystallized *in situ* via solvent evaporation. Mass spectra were acquired on a Bruker MicroFlex instrument operating in linear, positive-ion mode (25 kV accelerating voltage; 50-80% grid voltage), and final spectra were the result of averaging at least 50 laser shots. Two sets of molecular weight calibrants were used for low (4–20 kDa) and high (20–100 kDa) *m/z* ranges. Purification progress and sample MALDI spectra are illustrated in Supp Fig S1b and Fig 2, respectively.

**Figure 2.**
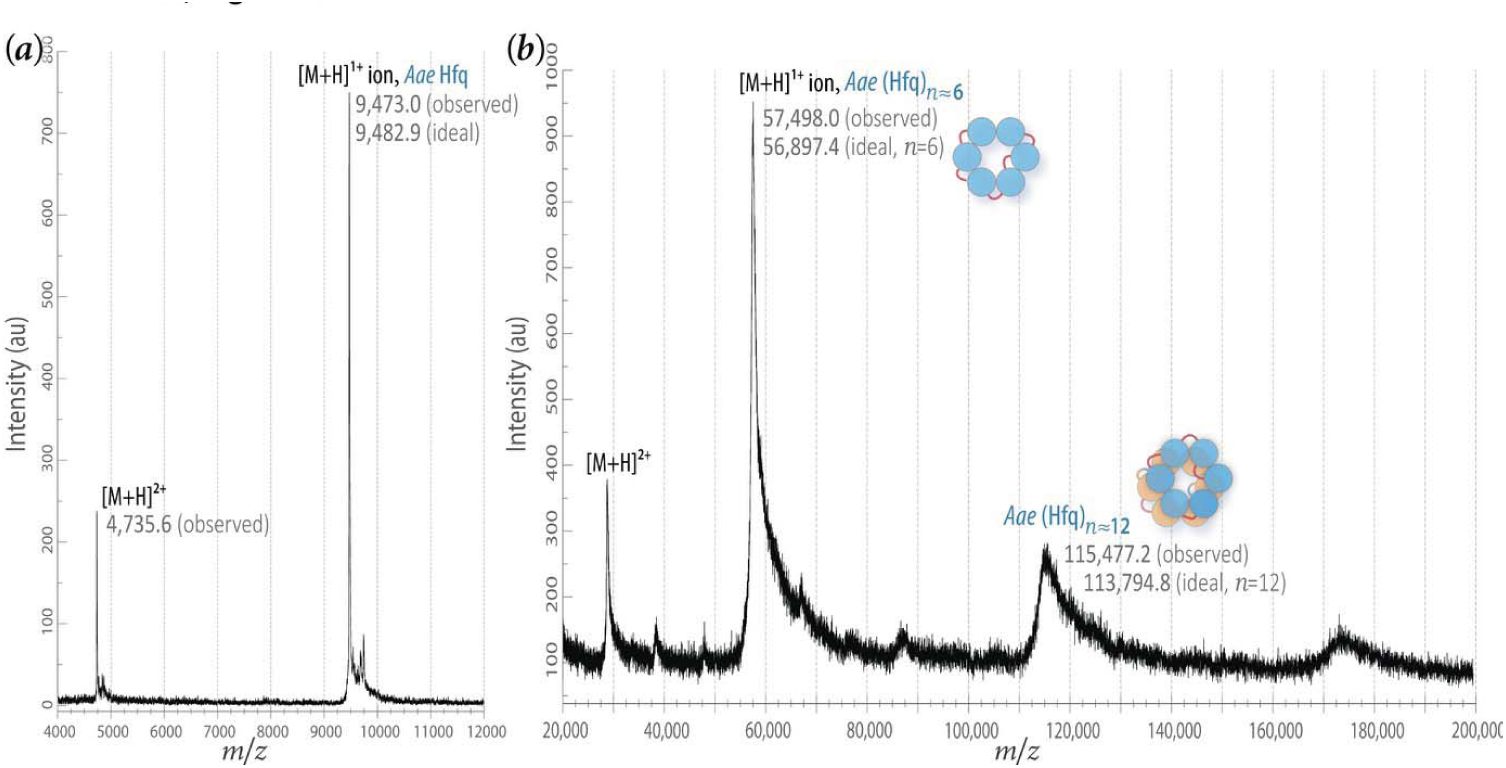
*Aae* Hfq monomers and oligomers, as assayed by crosslinking and mass spectrometry. MALDI-TOF spectra are shown for (*a*) native, untreated (non-crosslinked) *Aae* Hfq monomers, with an expected MW of 9482.9 Da based on the recombinant protein sequence (Supp Fig S1), as well as (*b*) a chemically crosslinked *Aae* Hfq sample. As detailed in the Methods (§2.2), crosslinking assays employed a gentle (‘indirect’) method, using formaldehyde as a crosslinking agent. The main peaks in the crosslinked sample correspond to hexamers and dodecamers, with expected MWs of 56,897.4 and 113,794.8 Da, respectively. The singly-charged molecular ion peaks, [M+H]^1+^, are accompanied by schematics (blue and orange balls) that indicate the anticipated architecture of the oligomeric states, alongside the peak’s MW, as determined from the mass spectrum (crosslinked species are better characterised by a MW range, rather than a single value, because of variability in the number of crosslinker molecules that react).

### 2.2. Crosslinking assays

Purified *Aae* Hfq was chemically crosslinked, using formaldehyde, in a so-called ‘indirect’ (vapor diffusion–based) method (Fadouloglou *et al.*, 2008). First, *Aae* Hfq samples at 0.6 mg/ml were dialyzed into a buffer consisting of 25 mM HEPES pH 8.0 and 500 mM NaCl. Reaction solutions were prepared in 24-well Linbro plates using micro-bridges (Hampton Research). Immediately before use, 5 N HCl was added to 25% w/v formaldehyde in a 1:40 v/v ratio. Next, 40 µl of this acidified formaldehyde solution was added to the micro-bridge, and 15 µl of the 0.6 mg/ml *Aae* Hfq was added to a silanized coverslip. Greased wells were then sealed by flipping over the coverslips and the reaction was incubated at 310 K for 40 min. Reactions were quenched by the addition of a primary amine; specifically, 5 µl of 1 M Tris pH 8.0 was mixed into the 15 µl protein droplet. Crosslinked samples were then desalted on a C4 resin (using *ZipTip*^®^ pipette tips) in preparation for analysis via MALDI-TOF MS as described above.

### 2.3. Analytical size-exclusion chromatography and multi-angle static light scattering

Analytical size-exclusion chromatography (AnSEC) was performed with a pre-packed *Superdex 200 Increase* 10/300 GL column and a Bio-Rad NGC™ medium-pressure liquid chromatography system. All protein samples were dialyzed into a running buffer consisting of 50 mM Tris pH 8.0 and 200 mM NaCl. *Aae* Hfq samples that were mixed with RNA sequences, denoted ‘U_6_’ (5′-monophosphate– r(U)_6_–3′-OH) or ‘A_18_’ (5′-monophosphate–r(A)_18_–3′-OH), were equilibrated by incubation at 310 K for 1 h prior to loading onto the AnSEC column. Elution volumes were measured by simultaneously monitoring the absorbance at 260 nm (RNA) and 280 nm (protein). A standard curve was generated using the Sigma gel-filtration markers kit, with calibrants in the 12–200 kDa molecular weight range: cytochrome c (12.4 kDa), carbonic anhydrase (29 kDa), bovine serum albumin (66 kDa), alcohol dehydrogenase (150 kDa) and β-amylase (200 kDa); blue dextran was used to calculate the void volume, *V*_0_.

To determine absolute molecular masses (i.e., without reference standards and implicit assumptions about spheroidal shapes), and in order to assess potential polydispersity of *Aae* Hfq in solution, multi-angle static light scattering (MALS) was used in tandem with size-exclusion chromatographic (SEC) separation. A flow-cell–equipped light scattering (LS) detector was used downstream of the SEC, in-line with an absorbance detector (UV) and a differential refractive index (RI) detector. In our SEC–UV/RI/LS system, (*i*) the SEC step serves to fractionate a potentially heterogeneous sample (giving the usual chromatogram, recorded at either 280 or 260 nm on a Waters UV/vis detector), (*ii*) the differential refractometer (RI) estimates the solute concentration via changes in solution refractive index (i.e., *dn*/*dc*), and (*iii*) the LS detector measures the excess scattered light. This workflow was executed on a Waters HPLC system equipped with the Wyatt instrumentation noted below, and utilized the same column and solution buffer conditions as described above (§2.1). LS measurements were taken at three detection angles, using a Wyatt miniDAWN TREOS (λ = 658 nm), and the differential refractive index was recorded from a Wyatt Optilab T-rEX. This enables the molecular mass of the solute in each fraction to be determined because the amount of light scattered (from the LS data) scales with the weight-averaged molecular masses (desired quantity) and solute concentrations (from the RI data); if multiple species exist in a given (heterogeneous) fraction, the polydispersity can be quantified as the ratio of the weight-averaged (M_*w*_) and number-averaged molar masses (M_*n*_). Data were processed and analysed using Wyatt’s ASTRA software package, applying the Zimm formalism to extract the weight-averaged molecular masses (Folta-Stogniew, 2009).

### 2.4. Fluorescence polarization–based binding assays

RNA–binding affinities were determined via fluorescence anisotropy/polarization experiments (FA/FP; (Pagano *et al.*, 2011)), using fluorescein-labelled oligoribonucleotides. In particular, the RNA probes 5′-FAM–r(U)_6_–3′-OH (FAM–U_6_) and 5′-FAM–r(A)_18_–3′-OH (FAM–A_18_) were used, with fluorescein amidite (FAM) modification of the 5′ ends; the FAM label features absorption and emission wavelengths, λ_*max*_, of 485 nm (excitation) and 520 nm (detection), respectively. FAM-labelled RNAs at 5 nM were added to a serially-diluted concentration series of purified *Aae* Hfq (in 50 mM Tris pH 8.0, 250 mM NaCl), and allowed to equilibrate for 45 minutes at room temperature. For binding assays that were supplemented with Mg^2+^, a 10 mM MgCl_2_ stock solution was used. The fluorescence polarization, *P*, is measured as 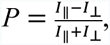 where *I*_||_ and *I*_⊥_ are the emitted light intensities in directions parallel and perpendicular to the excitation plane, respectively. FP data were recorded on a PheraSTAR spectrofluorimeter equipped with a plate reader (BMG Labtech), and values from three independent trials were averaged. The effective polarization, in units of millipolarization (*mP*), was plotted against log[(Hfq)_6_]. Binding data were fit, via nonlinear least-squares regression, to a logistic functional form of the classic sigmoidal curve for saturable binding. Specifically, the four-parameter equation

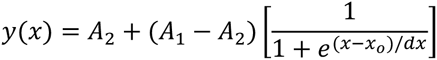

was used, where the independent variable *x* is the log[(Hfq)_6_] concentration at a given data point and the fit parameters are: (*i*) *A*_1_, the polarization at the start of the titration (unbound; lower plateau of the binding isotherm); (*ii*) *A*_2_, the final polarization at the end of the titration (saturated binding; upper plateau); (*iii*) *x*_0_, the apparent equilibrium dissociation constant (*K*_D_) for the binding reaction in terms of log[(Hfq)_6_]; and (*iv*) a parameter, *dx*, giving the characteristic scale/width over which the slope of the sigmoid changes (essentially the classic Hill coefficient, measuring the steepness of the binding curve). Calculations were performed with in-house code written in the R programming language, using the RSTUDIO integrated development environment.

### 2.5. X-ray crystallography

#### 2.5.1. Crystallization

Prior to crystallization trials, purified *Aae* Hfq was dialyzed into a buffer of 50 mM Tris pH 8.0 and 500 mM NaCl, and concentrated to 4.0 mg/ml. Protein samples were typically stored at 310 K, to retain solubility, and used within two weeks of purification. All crystallization trials were performed using the vapour diffusion method in sitting-drop format. Sparse-matrix screening (Jancarik & Kim, 1991) yielded initial leads (visible crystals) under several conditions, and these were then optimized by adjusting the concentration of protein and precipitating agent, as well as the pH of the mother liquor. Diffraction-grade crystals (Supp Figs S1c, d) were reproducibly obtained with 0.1 M sodium cacodylate pH 5.5, 5% w/v PEG-8000, and 40% v/v 2-methyl-2,4-pentanediol (MPD) as the crystallization buffer. In our final condition, 6-μl sitting drops (3 μl well + 3 μl of 4 mg/ml *Aae* Hfq) were equilibrated, at 295 K, against 600-μl wells containing the crystallization buffer. Initial micro-crystals developed over several days. Optimization of the above condition via additive screens (Hampton Research) led to the discovery of several compounds that, in a 1:4 v/v additive:crystallization buffer ratio, slowed nucleation and increased crystal size. The optimized crystals grew to average dimensions of 50x50x10 µm/edge within 2 weeks and adopted cubic or hexagonal plate morphologies. Three particularly useful additives, used in subsequent crystallization trials, were: (*i*) 0.1 M hexamminecobalt(III) chloride, [Co(NH_3_)_6_]Cl_3_, (*ii*) 1.0 M GndCl and (*iii*) 5% w/v of the non-ionic detergent *n*-octyl-β-D-glucoside. The final *apo*-form *Aae* Hfq crystals were obtained with additive (*i*); details are provided in Supp Table S2. *Aae* Hfq also was co-crystallized with a U-rich RNA (U_6_), under the above crystallization conditions and supplemented with additive (*ii*); these crystals were obtained by first incubating the purified protein with 500 µM 5′-monophosphate–r(U)_6_–3′-OH (hereafter denoted ‘U_6_’), in a 1:1 ratio, at 310 K for 1 h prior to setting-up the crystallization drop.

#### 2.5.2. Diffraction data collection and processing

The crystallization conditions described above adequately protected *Aae* Hfq crystals against ice formation upon flash-cooling (presumably because of the MPD), making it unnecessary to transfer crystals to an artificial mother liquor/cryo-protectant. Crystals were harvested using nylon loops and flash-cooled with liquid nitrogen. Diffraction data were collected at the Advanced Photon Source (APS) beamlines 24-ID-E and 24-ID-C for the *apo* and U_6_-bound crystal forms, respectively. Initial data-processing steps—indexing/integrating, scaling and merging—were performed in XDS (Kabsch, 2010). Space-group assignments and unit cell determinations utilized POINTLESS from the CCP4 suite (Winn *et al.*, 2011). Cell dimensions for the *apo* form (*P*1) are *a* = 63.45 Å, *b* = 66.06 Å, *c* = 66.10 Å, α = 66.05°, β = 83.94°, γ = 77.17°, and the U_6_–co-crystals (*P*6) have *a* = *b* = 66.19 Å, *c* = 34.21 Å.

#### 2.5.3. Structure solution, refinement and validation

Initial phases for the diffraction data-sets for both crystal forms were obtained via molecular replacement (MR). Specifically, the PHASER (McCoy *et al.*, 2007) software was used, with the *P. aeruginosa* (*Pae*) hexamer structure (PDB 1U1S) as a search model for phasing of both crystal forms (*Aae* and *Pae* Hfq share high sequence similarity; see Fig 1). Note that initial phases for the *P*1 and *P*6 *Aae* crystal forms were obtained independently of one another, *i.e.* via parallel MR efforts. For the *P*1 (*apo*) form, with 12 monomers/unit cell (indicative of two hexamers), the calculated Matthew’s coefficient (*V*_M_) is 2.06 Å^3^/Da, corresponding to a solvent content of 40.21% by volume. For the *P*6 (U_6_bound) form, only one monomer/ASU is feasible, with a *V*_M_ = 2.28 Å^3^/Da and a 46.08% solvent content. These and related characteristics are summarized in Table 1.

**Table 1.** X-ray diffraction data collection and processing statistics. Values for the highest-resolution shell are given in parentheses.

|  | *Aae* Hfq (*apo* form) | *Aae* Hfq•U<sub>6</sub> RNA |
| --- | --- | --- |
| Diffraction source | APS NE-CAT 24-ID-E | APS NE-CAT 24-ID-C |
| Wavelength (Å) | 0.97920 | 0.9195 |
| Temperature (K) | 100 | 100 |
| Detector | ADSC Q315 CCD | Dectris Pilatus 6MF |
| Crystal-detector distance (mm) | 200 | 300 |
| Rotation range per image (°) | 1.0 | 1.0 |
| Total rotation range (°) | 400.0 | 300.0 |
| Exposure time per image (s) | 1.0 | 1.0 |
| Space group | *P*1 | *P*6 |
| *a*, *b*, *c* (Å) | 63.46, 66.06, 66.10 | 66.19, 66.19, 34.21 |
| $\alpha$, $\beta$, $\gamma$ (°) | 60.05, 83.94, 77.17 | |
| Mosaicity (°) | 0.143 | 0.107 |
| Resolution range (Å) | 57.27–1.49 (1.53–1.49) | 34.21–1.50 (1.55–1.50) |
| Total No. of reflections | 299450 | 46203 |
| No. of unique reflections | 138120 | 13177 |
| Completeness (%) | 93.7 (83.7) | 94.9 (93.4) |
| Redundancy | 2.2 (2.1) | 3.5 (3.5) |
| $\langle I/\sigma(I) \rangle$ | 14.0 (3.4) | 12.3 (3.6) |
| $R_{\text{merge}}$ | 0.039 (0.258) | 0.056 (0.292) |
| $R_{\text{meas}}$ | 0.052 (0.349) | 0.065 (0.345) |
| $R_{\text{p.i.m.}}$ | 0.035 (0.234) | 0.032 (0.179) |
| $CC_{1/2}$ | 0.998 (0.886) | 0.998 (0.942) |
| Overall *B*-value from Wilson plot (Å<sup>2</sup>) | 12.62 | 15.87 |
| Matthews coefficient, $V_M$ (Å<sup>3</sup> Da<sup>−1</sup>) | 2.06 (for 12 subunits/AU) | 2.28 (for 1 subunit/AU) |
| Solvent content (% volume) | 40.21 | 46.08 |

<sup>†</sup> $R_{\text{sym}} = (\sum_{hkl} \alpha \sum_i |I_i(hkl) - \langle I_i(hkl) \rangle|) / (\sum_{hkl} \sum_i I_i(hkl))$ , where $I_i(hkl)$ is the intensity of the $i^{\text{th}}$ observation of reflection $hkl$ , $\langle \cdot \rangle$ denotes the mean of symmetry-related (or Friedel-related) reflections, and the coefficient $\alpha = 1$ ; the outer summations run over only unique $hkl$ with multiplicities greater than one.
<sup>‡</sup> $R_{\text{meas}}$ is defined analogously as $R_{\text{sym}}$ , save that the prefactor $\alpha = \sqrt{N_{hkl}/(N_{hkl} - 1)}$ is used; $N_{hkl}$ is the number of observations of reflection $hkl$ (index $i = 1 \rightarrow N_{hkl}$ ). Similarly, the precision-indicating merging *R*-factor, $R_{\text{p.i.m.}}$ , is defined as above but with the prefactor $\alpha = \sqrt{1/(N_{hkl} - 1)}$ .
<sup>§</sup> $CC_{1/2}$ is the correlation coefficient between intensities chosen from random halves of the full dataset.

After obtaining initial MR solutions in PHASER, the correct *Aae* Hfq amino acid sequence was built and side-chains completed in a largely automated manner, using the PHENIX suite’s AUTOBUILD functionality (Adams *et al.*, 2010). Individual solvent molecules, including H_2_O, MPD, and Gnd, were added in a semi-automated manner (i.e., with visual inspection and manual adjustment) after the initial stages of refinement. Refinement of atomic positions, occupancies and atomic displacement parameters (ADPs)—either as isotropic ‘*B*-factors’ or as full anisotropic ADPs—proceeded over several rounds in PHENIX. Some early refinement steps included simulated annealing torsion angle dynamics of coordinates, as well as refinement of TLS parameters to account for anisotropic disorder of each subunit chain (one TLS group defined per monomeric subunit). These steps yielded *R*_work_/*R*_free_ values of 0.194/0.212 and 0.212/0.223 for the *P*1 and *P*6 datasets, respectively. The diffraction limits of the *P*1 and *P*6 forms—1.49 Å and 1.50 Å, respectively—occupy an intermediate zone, between the atomic-resolution (*d* ≲ 1.4 Å) and medium resolution (*d* ≳ 1.7 Å) limits whereupon clearer decisions can be made as to the treatment of *B*-factors (Merritt, 2012). For instance, a relatively simple model (fewer parameters/atom), featuring individual isotropic *B*-factors and one TLS group per chain, might be most justifiable at ≈1.6 Å, depending on the quality of the diffraction data, whereas a more complex *B* model with a greater number of parameters—e.g., full anisotropic ADP tensors, *U*^*ij*^, one per atom—is likely to be statistically valid (and, indeed, advised) at resolutions better than ≈1.3 Å.

For both the *P*1 and *P*6 forms of *Aae* Hfq, a final *B*-factor model was chosen based on analyses of the data/parameter ratio (i.e., number of reflections/atom), Hamilton’s generalized residual (Hamilton, 1965) and related criteria, as implemented in the *bselect* routine of the PDB_REDO code (Joosten *et al.*, 2012). The *P*1 and *P*6 data-sets contained 16.5 and 17.5 reflections per atom, respectively, making the anisotropic refinement problem nearly two-fold overdetermined; PDB_REDO’s unsupervised decision algorithm identified the fully anisotropic, individual *B*-factor model as being optimal. The structural models resulting from various ADP refinement strategies were assessed using the protein anisotropic refinement validation and analysis tool (PARVATI; (Zucker *et al.*, 2010)). In the final refinement stages for both *Aae* Hfq crystal forms, *P*1 (*Z*=12 monomers/cell) and *P*6 (*Z*=6 monomers/cell), full anisotropic *B*-factor tensors were refined individually for virtually every atom. (A small fraction of atoms in both the *P*1 and *P*6 models were treated isotropically, i.e. by refining individual *B*_iso_ values; most of these atoms, selected based on per-atom statistical tests in PDB_REDO, were either water or heteroatoms [e.g., Gnd in *P*1, PEG in *P*6].) At no point in the refinement were NCS restraints or constraints imposed for the 12 subunits in the *P*1 cell. All steps of manual refinement and adjustment of the model were done in COOT (Emsley *et al.*, 2010).

After the correct protein sequence had been built and refined against the *P*6 dataset, at least two complete nucleotides of U_6_ RNA—including three phosphate groups—were clearly visible in σ_A_-weighted difference electron-density maps (*mF*_*o*_ − *DF*_*c*_). Ribonucleotides were built into electron density using the RCRANE utility (Keating & Pyle, 2010), after an initial round of refinement of coordinates, occupancies and individual *B*-factors in PHENIX. Validation of the final structural models included (*i*) inspection of the Ramachandran plot, via PROCHECK (Laskowski *et al.*, 1993); (*ii*) assessment of nonbonded interactions and geometric packing quality, via ERRAT (Colovos & Yeates, 1993); (*iii*) analysis of sequence/structure compatibility, via the profile–based method of VERIFY3D (Eisenberg *et al.*, 1997); and, finally, (*iv*) detailed stereochemical/quality checks with the MOLPROBITY software (Chen *et al.*, 2010). Final structure determination and model refinement statistics are provided in Table 2.

**Table 2.** Structure determination and model refinement. Values for the highest-resolution shell are given in parentheses.

|  | *Aae* Hfq (*apo* form) | *Aae* Hfq•U<sub>6</sub> RNA |
| --- | --- | --- |
| Resolution range (Å) | 46.35–1.49 (1.51–1.49) | 34.21–1.50 (1.56–1.50) |
| Completeness (%) | 93.9 | 94.9 |
| No. of reflections, working set | 138104 (12739) | 13171 (1308) |
| No. of reflections, test set | 10625 (983) | 662 (70) |
| Final $R_{\text{cryst}}$ | 0.1323 (0.1531) | 0.1443 (0.1499) |
| Final $R_{\text{free}}$ | 0.1696 (0.2108) | 0.1719 (0.1933) |
| No. of non-H atoms |  |  |
| Macromolecule | 7670 | 641 |
| Ligand | 267 | 15 |
| Solvent | 413 | 36 |
| Total | 8350 | 692 |
| R.m.s. deviations |  |  |
| Bonds (Å) | 0.005 | 0.005 |
| Angles (°) | 0.75 | 0.76 |
| Average $B$ factors (Å<sup>2</sup>) | | |
| Protein | 19.32 | 22.18 |
| Ligand | 25.89 | 30.44 |
| Ramachandran plot |  |  |
| Most favoured (%) | 98 | 97 |
| Allowed (%) | 1.7 | 2.9 |
| Outliers (%) | 0 | 0 |
| Rotamer outliers (%) | 0.34 | 1.5 |
| PDB ID | 5SZD | 5SZE |

### 2.6. Sequence and structure analyses

Sequences of verified Hfq homologs, drawn from diverse bacterial phyla, were selected for alignment and analysis against *Aae* Hfq. Here, we take ‘verified’ to mean that the putative Hfq homolog, from the published literature, has been identified via functional analysis or structural similarity (e.g., shown to adopt the Sm fold). Multiple sequence alignments were computed via two progressive alignment codes: (*i*) the multiple alignment using fast Fourier transform method (MAFFT; (Katoh & Standley, 2013)) and (*ii*) a sequence comparison approach using log-expectation scores for the profile function (muscle; (Edgar, 2004)). The geneious bioinformatics platform (Kearse *et al.*, 2012) was used for some data/project-management steps and tree visualization purposes. Multiple sequence alignments (Fig 1) were processed using ESPRIPT (Gouet *et al.*, 1999), run as a command-line tool; the resulting PostScript source was then modified to obtain final figures. Iterative PSI-BLAST (Camacho *et al.*, 2009) searches against sequences in the PDB were used to identify homologous proteins as trial MR search models. *Pae* Hfq, with 46% pairwise identity to *Aae* Hfq (across 97% query coverage), exhibited the greatest sequence similarity (≈63%, at the level of BLOSUM62) and was therefore chosen as the initial MR search model.

Structural alignments were performed using a least-squares fitting algorithm (McLachlan, 1982) implemented in the program PROFIT (Martin & Porter, 2009). Multiple structural alignment of the 12 monomeric subunits in the *apo* form of *Aae* Hfq was used to create a mean reference structure, and each monomer was then aligned to that averaged reference. To assess 3D structural similarity between each of the *n*(*n–*1)⁄2 distinct pairs of monomers, a pairwise distance matrix was constructed by computing main-chain RMSDs between subunits *i* and *j*, giving matrix element (*i*, *j*). Agglomerative hierarchical clustering was performed on this distance matrix, using either the complete–linkage criterion or Ward’s variance minimization algorithm with a Euclidean distance metric (Jain *et al.*, 1999); in-house code was written for these steps in both the R (within RSTUDIO) and Python languages.

Secondary structural element residue boundaries were determined by a consensus approach, via visual inspection in PYMOL as well as the automated assignment tools DSSP and STRIDE. Normal mode analyses of the *P*1 and *P*6 structures—taken as coarse-grained (C_α_-only) representations and treated as anisotropic network models (ANM)—were performed with the PRODY/NMWIZ (Bakan *et al.*, 2011) plugin to VMD (Humphrey *et al.*, 1996), using default parameters for ANM spring constants and interaction cutoff distances. Of the 3*N*–6 nontrivial modes, displacements along the softest ≈ 20 vibrational modes, which correspond to low-frequency/high-amplitude collective motions, were visually inspected in VMD. Other structure analyses (e.g., Fig 6a) entailed computing the principal axes of the moment of inertia tensor and the best-fit plane to 3D structures (in the sense of linear least-squares); the latter task utilized a previously-described singular value decomposition code (Mura *et al.*, 2010), and all other structural analysis tasks employed in-house code written in Python or as Unix shell scripts. Nucleic acid stereochemical parameters and conformational properties, e.g. values of glycosidic torsion angles and sugar pucker phase angles of the U_6_ RNA, were analysed and calculated with the program DSSR (Lu *et al.*, 2015). Surface area properties, such as solvent-accessible surface areas (SASA) and buried surface areas (BSA, or ΔSASA), were calculated as averages from five approaches: (*i*) Shrake & Rupley’s ‘surface-dot’ counting method (Shrake & Rupley, 1973), as implemented in AREAIMOL; (*ii*) the classic Lee & Richards ‘rolling-ball’ method (Lee & Richards, 1971), available in NACCESS; (*iii*) the ‘reduced surface’ analytical approach of MSMS (Sanner *et al.*, 1996); and more approximate (point-counting) methods from the structural analysis routines available in (*iv*) PyMOL and (*v*) PyCogent (Cieślik *et al.*, 2011).

All molecular graphics illustrations in Figs 5–8 and Supp Figs S3–S6 were created in PyMOL, with the exception of Fig S4e, f (created in VMD, rendered with Tachyon). LigPlot+ (Laskowski & Swindells, 2011) was used in creating schematic diagrams of interatomic contacts, as in Fig 8. Many of our scientific software tools were used as SBgrid–supported applications (Morin *et al.*, 2013).

## 3. Results

The organism *A. aeolicus* belongs to the taxonomic order *Aquificales*, in the phylum *Aquificae*, within what may be the most phylogenetically ancient and deeply branching lineage of the *Bacteria*. Thus, this species offers a potentially informative context in which to examine the evolution of sRNA-based regulatory systems, such as those built upon Hfq. The *Aae* genome contains an open reading frame with detectable sequence similarity to characterized Hfq homologs (e.g., from *E. coli* and other proteobateria), and an RNomics/deep-sequencing study has shown that this putative Hfq homolog, upon heterologous expression in the γ-proteobacterium *Salmonella enterica*, can immunoprecipitate host sRNAs (Sittka *et al.*, 2009). Sequence analysis confirms that this putative Hfq can be identified via database searches (Fig 1), and that this homolog exhibits enhanced residue conservation at sequence positions that correspond to the three RNA-binding sites on the surface of Hfq (*proximal*, *distal* and the *lateral rim*, denoted in the consensus line in Fig 1). As the first step in our crystallographic studies, we cloned, expressed and purified recombinant *Aae* Hfq; in these initial experiments, *Aae* Hfq generally resembled hitherto characterized Hfq homologs in terms of biochemical properties (e.g., resistance to chemical and thermal denaturation, hexamer formation).

### 3.1. Cloning, expression, purification and initial biochemical examination of *Aae* Hfq

Recombinant, wild-type *Aae* Hfq was successfully cloned, over-expressed and purified from *E. coli*, as confirmed by various biochemical and biophysical data, including SDS-PAGE gels (Supp Fig S1) and MALDI-TOF mass spectra of the native protein (Fig 2a). The His6×-tagged *Aae* Hfq is 100 amino acids (AA) long, with a molecular weight of 11,365.0 Da and a predicted isoelectric point of 9.69; the working *Aae* Hfq construct, obtained via proteolytic removal of the tag (Supp Fig S1a), is 83 AA (9,482.9 Da, pI = 9.45). The expected mass, computed from the AA sequence, is in close agreement with that experimentally characterized by MALDI–TOF, indicating successful (complete) removal of the affinity tag (Fig 2a) at position G^−2^ (residue numbering is such that the native methionine is M^1^).

Initial *Aae* Hfq purification efforts were hindered by nucleic acid contaminants. Specifically, purified protein samples exhibited A_260_/A_280_ absorbance ratios of ≈ 1.65, indicative of co-purifying nucleic acids (De Mey *et al.*, 2006, Patterson & Mura, 2013); this problem is perhaps unsurprising, given the known affinity of Hfq for nucleic acids, combined with *Aae* Hfq’s particularly high pI. By applying systematic colorimetric assays (Patterson & Mura, 2013) to *Aae* Hfq samples with high A_260_/A_280_ ratios (Supp Fig S2a), we found that the co-purifying nucleic acids likely comprise a heterogeneous pool of RNAs, with lengths between ≈ 100-200 base pairs (Supp Fig S2b). Early experiments using anion-exchange chromatography revealed that nucleic acid–bound Hfq would elute at three distinct ionic strengths (in a linear salt gradient), and each peak appeared to contain a population of nucleic acids that varied in length, both within one peak and between the three peaks (data not shown). To obtain well-defined, well-behaved *apo Aae* Hfq samples—for downstream RNA-binding assays, crystallization trials, etc.—relatively high concentrations (≈ 6 M) of guanidinium were added to cell lysates, the aim being to dissociate spurious Hfq-associated nucleic acids. Inclusion of Gnd in the purification workflow (see Methods) yielded samples with improved A_260_/A_280_ ratios (≈ 0.8), suggesting that nucleic acid contamination had been alleviated. Notably, the Gnd denaturant did not appear to unfold or disrupt *Aae* Hfq’s oligomerization properties, based on various observations; for instance, a discrete band corresponding to the hexameric assembly persisted on SDS-PAGE gels of Gnd-treated samples (Supp Fig S1b).

As an initial assessment of its self-assembly properties and oligomeric states in solution, purified *Aae* Hfq was examined by analytical size-exclusion chromatography (Fig 3a, b black traces). The protein elutes as a single, well-shaped peak, with no apparent splitting, broadening, shouldering, tailing, etc. However, the location of this peak is unexpected: the peak elution volume gives a molecular weight (MW) of ≈ 37 kDa, rather than the ≈ 57 kDa expected for an *Aae* Hfq hexamer. This apparent MW, obtained using a standard curve as described in the Methods section, could indicate a tetrameric assembly, for which the MW is calculated to be 37.9 kDa. Shape-dependent deviations from ideal migration properties would be expected to give an (Hfq)_6_ species that migrates *faster*, not slower, than anticipated based purely on MW, given the larger effective hydrodynamic radius of a toroidal hexamer (versus the roughly globular standards used to calibrate our column elution volumes). However, favourable protein⋯resin interactions would tend to retard the migration of an *Aae* Hfq oligomer, leading to a smaller apparent MW species. Given the highly basic pI, and resultant charge on *Aae* Hfq at near-neutral pHs, we suspect that the low MW estimate from AnSEC stems from protein⋯resin interactions, electrostatic or otherwise; spurious *Aae* Hfq retention was also seen in experiments with other, unrelated chromatographic resins; note that nonspecific protein adsorption to SEC resins was first documented long ago (Belew *et al.*, 1978) and has been reviewed (Arakawa *et al.*, 2010).

**Figure 3.**
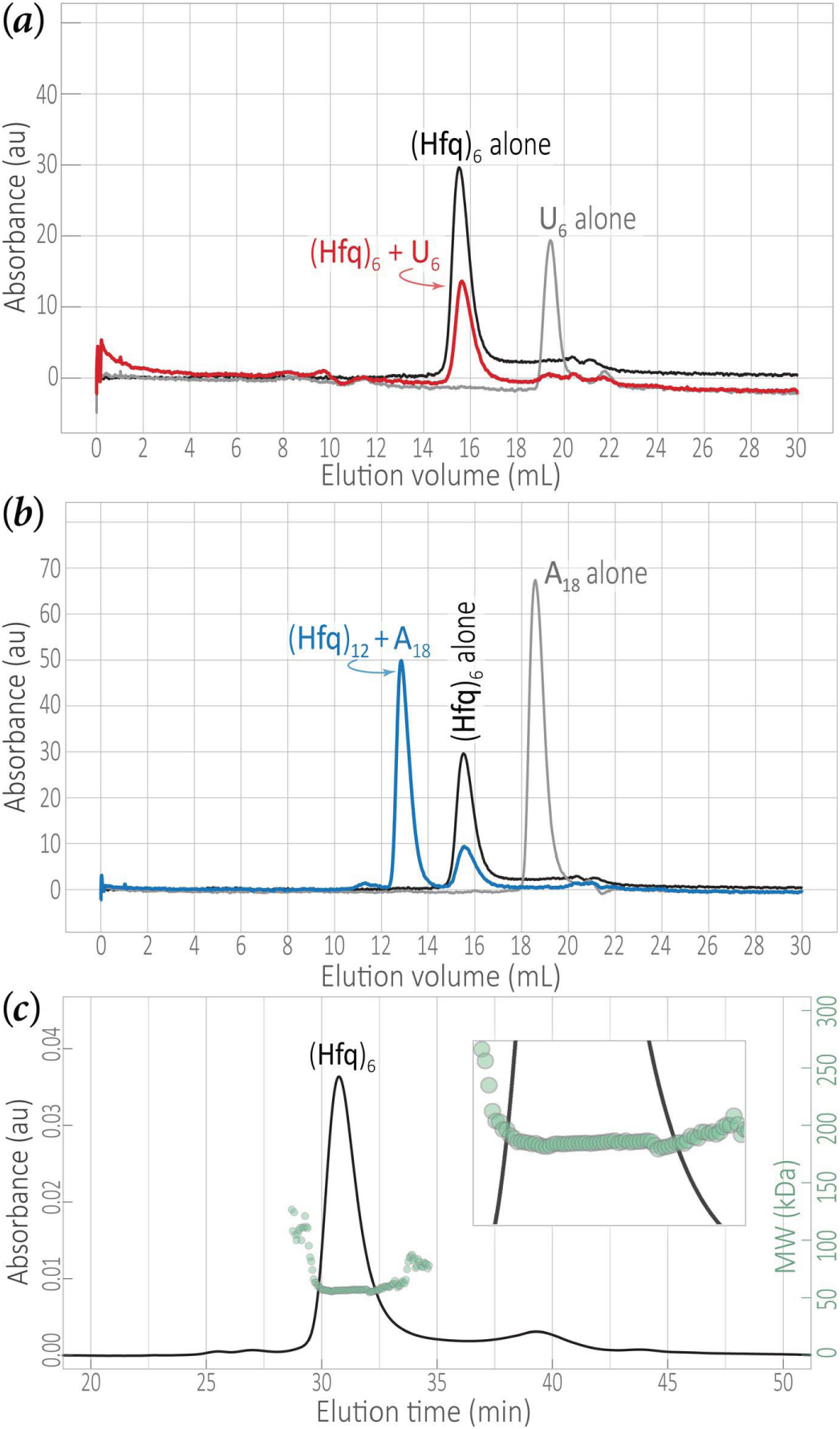
The solution-state distribution of *Aae* Hfq oligomers shifts in the presence of short RNAs. Elution profiles are shown for analytical size-exclusion chromatography of *Aae* Hfq samples incubated with either (*a*) U_6_ or (*b*) A_18_ RNAs. The elution of *Aae* Hfq was detected via the absorbance at 280 nm (A_280_), and RNA and Hfq•RNA complexes were monitored at A_260_. While putative Hfq⋯U_6_ interactions do not appear to shift the oligomeric state, as indicated by the close alignment of the black (Hfq alone) and red (Hfq•U_6_) peaks in (*a*), Hfq interactions with A_18_ do shift the oligomeric species towards a higher-order state (blue arrow in *b*, denoting apparent dodecamers). This shift could correspond to the simultaneous binding of A_18_ to two Hfq hexamers, potentially via two modes: (*i*) as an (Hfq)_6_•A_18_•(Hfq)_6_ ‘bridged’ complex, or (*ii*) as A_18_ bound to one of the two distal faces that would be exposed on an independently-stable (Hfq_6_)_2_ double-ring dodecamer. These two models cannot be distinguished via AnSEC. (*c*) To verify the molecular weight of the *Aae* Hfq elution peak, the protein was analysed via SEC fractionation followed by multi-angle static light scattering and refractive index measurements. The SEC elution profile (black trace) is taken as the absorbance at 280 nm. Light-scattering and refractive index data can be used to compute molar masses, and the open circles shown here (semi-transparent green) are the molar mass distribution data (i.e., masses [in kDa] as a function of elution volume). The weight-averaged molecular weight, M_w_, of the Hfq sample is computed for the entire peak from this distribution, and the scale is given by the vertical axis on the right-hand side (green numbers; note that this scale applies to the main plot, not the inset). The apparent M_w_ that was computed, 58.75 kDa, corresponds to a hexameric assembly of *Aae* Hfq.

The aberrant AnSEC elution behaviour prompted us to assay the *Aae* oligomeric state by alternative means. SEC coupled with multi-angle light scattering (MALS) showed that the *Aae* Hfq eluting at this peak position corresponds to a hexamer, with a weight-averaged molecular weight, M_w_, of 58.75 kDa (Fig 3c). A plot of the molar mass distribution (Fig 3c, green circles) exhibits uniform values across this *Aae* Hfq peak (Fig 3c, inset), indicating that this region of the eluted sample is mono-disperse. *Aae* Hfq monomers were susceptible to chemical crosslinking with formaldehyde, as analysed by MALDI–TOF MS (Fig 2). The main peak in the mass spectrum of this sample (Fig 2b) corresponds to a hexamer (57,498.0 Da from MS, versus 56,897.4 Da from the sequence); a second peak, near ≈ 115 kDa, corresponds to within 1.5% of the MW of a dodecameric assembly. Some Sm and Hfq orthologues have been found to assemble into stacked double-rings and other higher-order species, based on analytical ultracentrifugation and light-scattering data (Mura, Kozhukhovsky*, et al.*, 2003, Mura, Phillips*, et al.*, 2003, Dimastrogiovanni *et al.*, 2014), electron microscopy (Arluison *et al.*, 2006, Mura, Kozhukhovsky*, et al.*, 2003), gel-shift assays and other approaches; however, an integrated experimental analysis, using multiple independent methodologies on the same Hfq system, strongly suggests that the (Hfq)_6_•RNA binding stoichiometry is predominantly 1:1 (Updegrove *et al.*, 2011).

### 3.2. Characterization of RNA-binding by *Aae* Hfq in solution

To evaluate putative RNA interactions with *Aae* Hfq, solution-state binding interactions between *Aae* Hfq and either U_6_ or A_18_ (unlabelled) RNAs were examined via analytical size-exclusion chromatography. RNAs that are U-rich (e.g., U_6_) or A-rich (e.g., harbouring an (AAN)_*n*_ motif) are known to bind at the proximal and distal faces, respectively, of Hfq homologs from Gram-negative species. U_6_ RNA was shown to bind *Aae* Hfq in solution via comparison of the following elution profiles (Fig 3a): (*i*) Hfq-only (black trace, detected via absorbance at 280 nm), (*ii*) U_6_-only (grey, monitored at 260 nm) and (*iii*) an Hfq+U_6_ mixture (red, 260 nm). In sample (*iii*), the Hfq+U_6_ mixture, note the absence of a U_6_ RNA peak near 19.5 mL (Fig 3a, grey), and concomitant peak shift centred at the Hfq-only trace, indicative of saturated binding of the RNA. Properties of the elution profiles for samples (*i*) and (*iii*)—specifically, no shift in the peak position and no alteration of the bilateral symmetry of the peak—suggest that the addition of U_6_ does not alter the apparent monomer↔hexamer equilibrium of *Aae* Hfq.

In contrast to the U_6_ behaviour, adding A_18_ RNA to an *Aae* Hfq sample does appear to shift the Hfq oligomeric state to a higher-order species (Fig 3b, blue trace, major peak) that coexists with the usual hexamer (blue trace, minor peak). This newly-appearing, A_18_-induced species is hydrodynamically larger than (Hfq)_6_, as it elutes far earlier than does Hfq in the Hfq-only sample (black trace); the higher-order appears to correspond to an *Aae* Hfq dodecamer. This was further verified based on the M_*w*_ determined through SEC-MALS experiments done in parallel (data not shown). Also, note that the Hfq+A_18_ trace is devoid of a peak at the A_18_-only position (i.e., no peak in the blue trace, near the ≈ 18.5 mL peak location of the grey trace), indicating that binding has saturated with respect to A_18_.

To further quantify the interactions of Hfq with U-rich and A-rich RNAs, the binding affinities for *Aae* Hfq with 5′-FAM–labelled RNA oligoribonucleotides were determined via fluorescence polarization (FP) assays (Fig 4). We took FAM-U_6_ and FAM-A_18_ probes as proxies for U-rich and A-rich ssRNAs, enabling us to assay the strength of *Aae* Hfq⋯RNA interactions with these prototypical A/U-rich RNAs (for brevity, we simply refer to these RNAs as ‘U_6_’ and ‘A_18_’ if the FAM is obvious from context). Both U_6_ and A_18_ were found to bind *Aae* Hfq with similarly high affinities: the nanomolar–scale dissociation constants (*K*_D_) are 21.3 nM for U_6_ and 17.4 nM for A_18_ (Fig 4, thin traces). The inclusion of 10 mM Mg^2+^ in the binding reaction enhanced the U_6_–binding affinity by an order of magnitude, yielding a *K*_D_ of 2.1 nM (Fig 4; red, thicker trace); the A_18_–binding affinity also increased in the presence of Mg^2+^, though by only two-fold, to a *K*_D_ of 9.5 nM (blue, thicker trace). No significant binding was detected between *Aae* Hfq and either FAM-A_6_ or FAM-C_6_ (data not shown).

**Figure 4.**
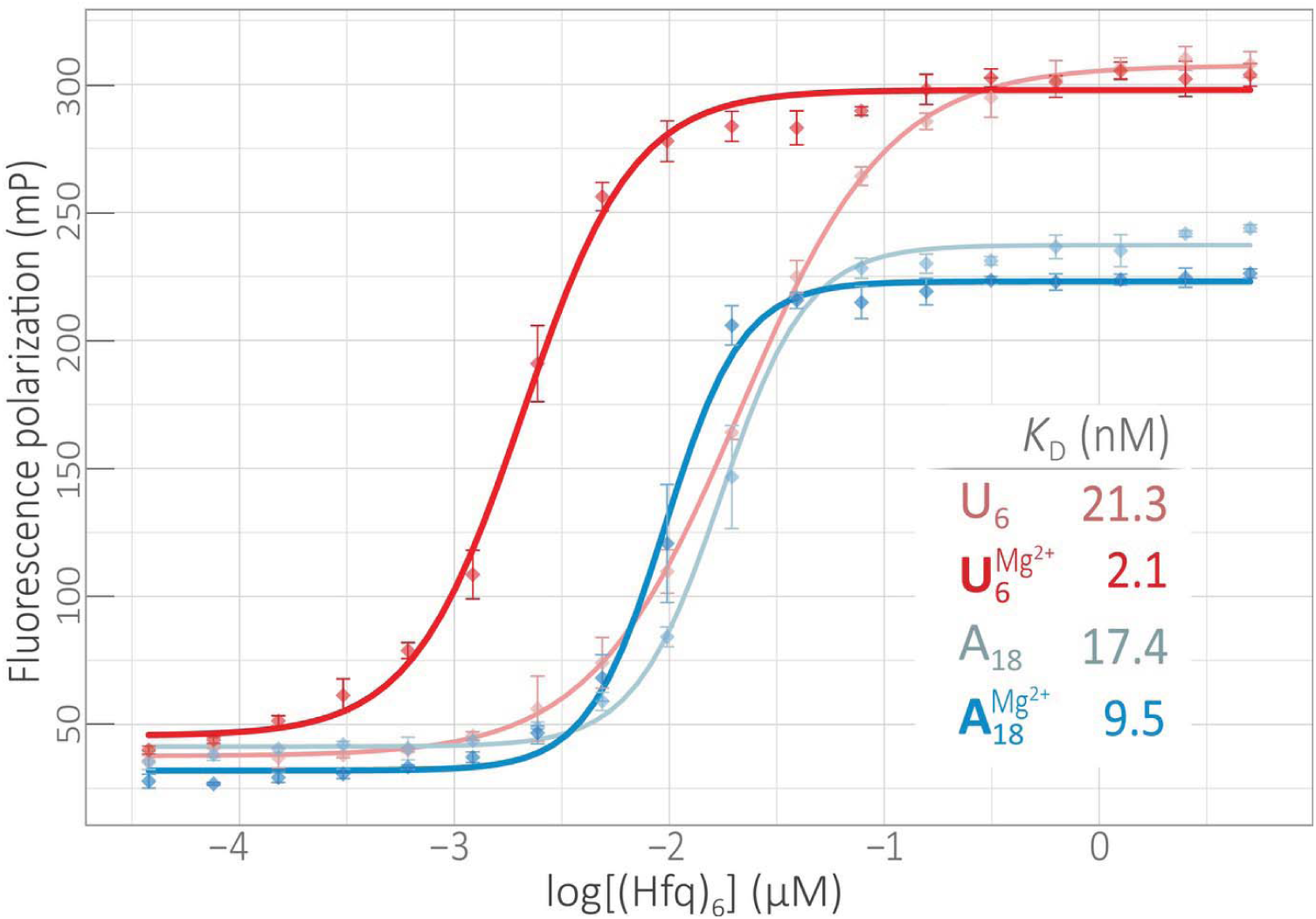
High-affinity binding of *Aae* Hfq to A– and U–rich RNAs, with variable Mg^2+^ dependencies. Binding was quantified via fluorescence polarization assays using 5 nM FAM-U_6_ (red) or FAM-A_18_ (blue) and varying concentrations of Hfq, either in the absence (thin lines) or presence (thick lines) the addition of 10 mM MgCl_2_. For each binding reaction, data from three replicates (standard errors given by vertical bars) were fit using a standard, sigmoidal binding isotherm equation (§2.4). The computed binding constants are given (inset) in terms of the (Hfq)_6_ concentration (the stoichiometry of all characterized Hfq•RNA complexes, as well as the structural results reported herein, suggest that the hexamer is the active/functional unit). The addition of Mg^2+^ increases the binding affinity for both FAM-U_6_ and FAM-A_18_, albeit with a greater influence for the U-rich (proximal site–binding) RNA. Significant binding was not detected for a shorter A-rich (FAM-A_6_) or C-rich (FAM-C_6_) ssRNA.

### 3.3. Crystal structures of *Aae* Hfq monomers and oligomers, and the lattice packing

Crystals of *Aae* Hfq were readily obtained in multiple forms, including hexagonal plates and small, birefringent parallelepiped habits (Supp Fig S1c). At least three distinct morphologies could be identified, which we denote (*i*) a ‘*P*1 form’ (*apo* Hfq, without RNA), (*ii*) a ‘*P*6 form’ (with RNA, see §3.4 below) and (*iii*) a third form that likely belongs to space-group *P*3_1_ or *P*6_2_. Forms (*i*) and (*ii*) were well-diffracting (Supp Fig S1d), leading to the *P*1 and *P*6 structures reported here; the third form yielded diffraction data with potential pathologies, including translational pseudosymmetry or tetartohedral twinning, and its structure is the subject of future work (Stanek & Mura, unpublished data). Initial *Aae* Hfq crystals were obtained with a crystallization reagent comprised of 0.1 M sodium cacodylate, 5% w/v PEG 8000 and 40% v/v MPD; inclusion of [Co(NH_3_)_6_]Cl_3_ additive, at ≈10 mM in the final crystallization drop, improved specimen size and quality. These *apo Aae* Hfq crystals formed in space-group *P*1, with cell dimensions *a* = 63.46 Å, *b* = 66.06 Å, *c* = 66.10 Å, α = 60.05°, β = 83.94°, γ = 77.17°. These dimensions are most consistent with *Z* ≈ 10–12 monomers/cell, and a resolution-dependent probabilistic estimator for the Matthews coefficient (Kantardjieff & Rupp, 2003) gives a 12-mer as the second highest peak; also, the *a* ≈ *b* ≈ *c* geometry is consistent with a model wherein two Hfq hexameric rings, which generally measure ≈ 65 Å in diameter, stack atop one another in the cell.

The *P*1 *Aae* Hfq structure was refined to 1.49 Å resolution, with initial phases obtained by molecular replacement with a *Pae* Hfq hexamer search model (PDB 1U1S; (Nikulin *et al.*, 2005)). The *Pae* homolog was used because sequence analysis (Fig 1) showed it to have the greatest sequence identity (>40%) to *Aae* Hfq. A promising molecular replacement solution was readily identified, and side-chains for the *Aae* Hfq sequence were initially built in an automated manner using PHENIX. As detailed in the Methods section, the number of reflections per atom, as well as other diffraction data quality statistics, prompted us to refine atomic displacement parameters (ADPs) via treatment of the full, anisotropic *B*-factor tensor for essentially all non-hydrogen atoms (most of the isotropically-treated exceptions were atoms of solvent molecules or small-molecule components of the crystallization buffer). Anisotropic treatment of individual ADPs began at a relatively late stage in the overall refinement workflow, and doing so noticeably improved the *R*_work_/*R*_free_ residuals, from 13.6%/17.2% to 12.8%/15.6% before and after anisotropic treatment, respectively (Table 2). The final, refined *P*1 model was subjected to extensive validation and quality assessment, as described in the Methods section, in terms of both the 3D structure itself (i.e., atomic coordinates) as well as the patterns of *B*-factors (i.e., anisotropic ADPs).

The overall 3D structure of the *Aae* Hfq monomer (Fig 5) is that of the Sm fold, as anticipated based on sequence similarity and the efficacy of MR in phasing the diffraction data. In particular, an *N*-terminal α-helix is followed by five highly-curved β-strands arranged as an antiparallel β-sheet. The secondary structural elements (SSEs), shown schematically in Fig 1, are labelled in the 3D structure of Fig 6b. Precise SSE boundaries in *Aae* Hfq, computed with STRIDE, are residues # 5–16 (α1), 19–24 (β1), 29–38 (β2), 41–46 (β3), 49–54 (β4) and 58–63 (β5); the same ranges are obtained with DSSP, save that DSSP’s criteria make F37 (not D38) the end of the most curved strand (β2). Most of *Aae* Hfq’s β-strands are delimited by loops that adopt various β-turn geometries (including types I, II′, IV, VIII), with the exception of a short 3_10_ helix (residues 55–57) between β4→β5. These loops contain many of the RNA-contacting residues of Hfq (see below) and, as labelled in Figs 1, 5, 6 and 9, we denote these linker regions as L1→5. Noncovalent interactions between Hfq monomers include van der Waals contacts and hydrogen bonds between the backbones of strand β4 of one subunit and β5* of the adjacent subunit, effectively extending the β-sheet across the entire toroid; these enthalpically favourable interatomic contacts likely facilitate self-assembly of the hexamer. (Unless otherwise stated, asterisks denote an adjacent Hfq subunit, be it related by crystallographic symmetry or otherwise.) Residues 1→68 of the native *Aae* Hfq sequence could be readily built into electron density maps for each monomer in the ASU, thus providing a structure of Hfq’s *N*-terminal region as well as the entire Sm domain; note that the *N*-terminal tail, illustrated for the *apo/P*1 structure in Fig 5 (bottom-right) and Fig 6b, is unresolved in many Hfq structures. Most of the *C*-terminal residues 70→80 were not discernible in electron density, and are presumably disordered.

**Figure 5.**
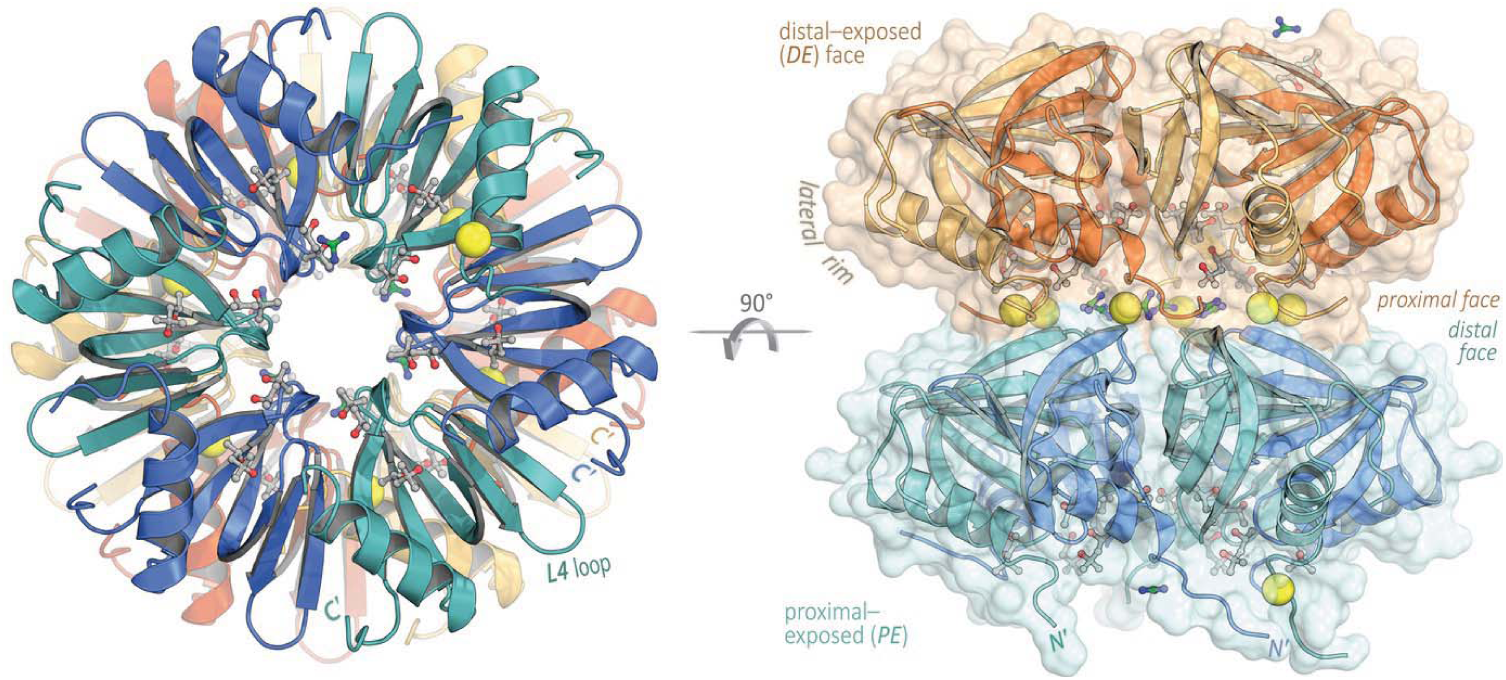
Crystal structure of *Aae* Hfq in the *apo* form, with head–to–tail stacking of hexameric rings. The *apo* form of *Aae* Hfq crystallized in *P*1 as a dodecameric assembly of hexamers, stacked in a *proximal*-to-*distal* orientation in the lattice. Ribbon diagrams of the final, refined structure are shown here, from perpendicular viewpoints. The proximal-exposed (*PE*) hexamer is coloured blue and cyan, and subunits in the distal-exposed (*DE*) hexamer are coloured alternatingly yellow and orange. Co-crystallizing molecules of MPD (grey carbons) and GndCl (green carbons) are shown as ball-and-stick representations, and Cl^−^ ions are rendered as yellow spheres scaled to the van der Waals radius. Note that many of the Gnd cations and Cl^−^ anions are coplanar, where they form a ‘salty’ layer at the ring interface (this is most clearly seen in the transverse view). Contacts between hexamers are mediated by the *N*-termini of the *DE* hexamer (top) and the loop L2/strand β2 regions of the *PE* hexamer (bottom); the approximate location of one of the lateral RNA-binding sites is labelled on the *DE* ring.

**Figure 6.**
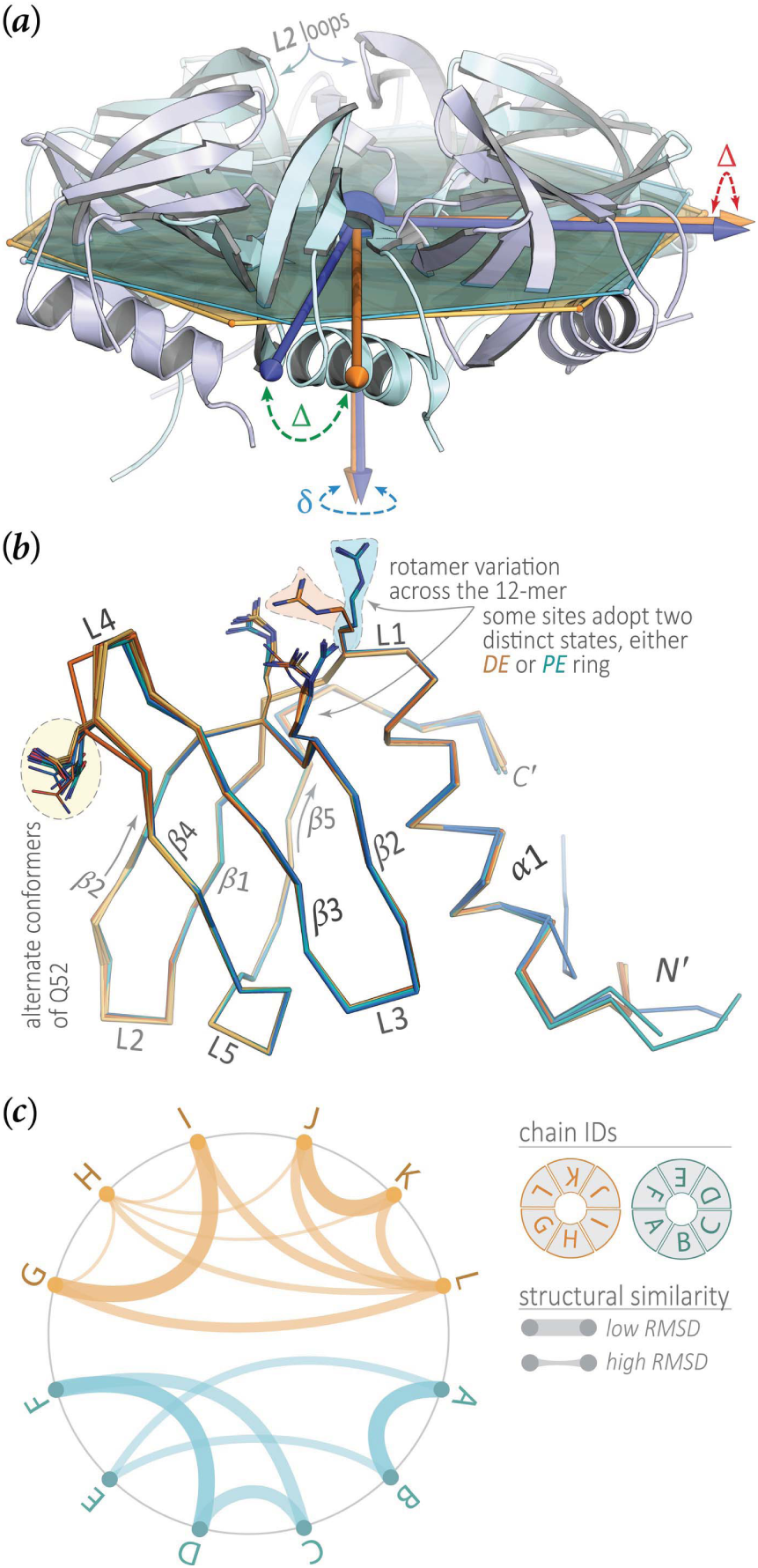
Structural variation across the *Aae* Hfq monomer (*P*6) and dodecamer (*P*1) crystal forms. At a gross structural level, the two Hfq rings in the head-tail dodecamer of the *P*1 crystal form (Fig 5, axial view) appear to be related by a rigid-body rotation. The two rings—the proximal-exposed (*PE*) and distal-exposed (*DE*) hexamers—were brought, via pure rigid-body translation, to a common origin, indicated by the blue sphere in (*a*). Best-fit planes to each ring were then computed, as described in the Methods section (§2.6) and shown here as semi-transparent hexagonal plates of either orange (*DE* ring) or cyan (*PE* ring) colour. For clarity, the *DE* ring (orange/yellow in Fig 5) is omitted in panel (*a*), and a couple of the L2 loops are labelled (in the *PE* ring) simply as a structural landmark. The three principal axes of the moment of inertia tensor are shown in either orange (*DE* ring) or blue (*PE* ring); large differences in the orientation of these principal axes are marked by green and red ‘Δ’ symbols, while a ‘δ’ symbol (blue) denotes smaller-scale differences. The rotation between the rings is clear from the relative disposition (Δ) of two of the principal axes. Furthermore, a small— but discernable—difference (δ) in the directions of the normal axes indicates a slight tilt between the rings; this direction would correspond to the 6-fold axis in a perfectly symmetric hexamer. A multiple structural alignment of the 12 subunits in the *P*1 cell (*b*) reveals little structural variation of the Sm core (shown as C_α_ backbone traces), while there are many examples of side-chain variability (as noted in the panel). The defining secondary structural elements of the Sm fold (L1 loop, β1 strand, etc.), as well as the termini, are labelled. The two regions of *Aae* Hfq that most extensively engage in interactions between rings (*hexamer⋯hexamer* contacts in Fig 5), and in forming crystal contacts, are the L4 loops and the irregularly-structured ≈5 residues at the *N*-terminus (preceding α1). These also are the two most variable regions in Hfq, both in terms of sequence length (and composition) as well as 3D structure, as seen in (*b*). The side-chain variability shown in (*b*) takes two forms: (*i*) alternate conformers that could be built for a single residue, such as the Q52 example highlighted to the left, and (ii) rotameric variation for a single residue across the 12 subunits, such as the groups of three residues shown as sticks near the top of (*b*). In many instances of the latter case, the 12 residue states clustered into two groups, corresponding to the *DE* or *PE* hexamer. In the diagram of panel (*c*), the Hfq subunits in *P*1, labelled by chain ID, are evenly spaced about a circle; arcs are drawn between the most structurally similar pairs of subunits, with the line thickness inversely scaled by the RMSD for the given pair. For clarity, not all ≈ *n*^2^ edges are shown here, but rather only at the levels of subunit pairs and triples (i.e., the deepest and second-deepest levels of leaf-nodes in the full dendrogram of Supp Fig S3c). This result, from hierarchical clustering on backbone RMSDs, shows that pairs of monomers within a given hexamer are structurally more similar to each other than are pairs between hexamers (chains A→F comprise the *PE* ring and G→L are the *DE* ring).

While neither NCS averaging, nor any NCS constraints or restraints, were applied at any point in the phasing and refinement of *Aae* Hfq in the *apo* form, the 12 monomers in the *P*1 cell are virtually indistinguishable from one another (Fig 6a,b, Supp Fig S3), at least at the level of protein backbone structure (there are side-chain variations). The mean pairwise main-chain RMSD, for all monomer pairs in the *P*1 cell, lies below 0.3 Å; this low value is also evident in the magnitude of the ordinate scale of the structural clustering dendrogram in Supp Fig S3c. To systematically compare structures, a matrix of RMSDs was constructed from all pairwise subunit alignments. Agglomerative hierarchical clustering on this distance matrix (Supp Fig S3c) reveals that the subunits partition into two low-level (root-level) clusters so as to recapitulate the natural (structural) ordering in the crystal: that is, chains A→F cluster together (as the *proximal-exposed*, or *PE*, ring in Fig 5), and likewise chains G→L form a second group (the *distal-exposed*, or *DE*, ring). This finding is illustrated in Fig 6c, which conveys the degree of 3D structural similarity as a circular graph wherein an edge between two chains is inversely scaled by their RMSD.

At the *Aae* Hfq monomer level, the greatest structural variation occurs among the *N*-termini and the L4 loop region between β3→β4; apart from the termini, loop L4 (Fig 6b) is the most variable region in most known protein structures from the Sm superfamily. The conformational heterogeneity in the termini and loops of *Aae* Hfq stems, at least partly, from differing patterns of interatomic contacts for different subunits, at the levels of monomers, hexamers and dodecamers in the overall *P*1 lattice. The patterns of conformational heterogeneity are clear when the dodecameric structure is visualized as a cartoon, with the diameter of the backbone tube scaled by the magnitude of per-atom *B*_eq_ values (this derived quantity, computed from the trace of the full anisotropic ADP tensor, is taken as an estimate of the true *B*_iso_ values that would result from refinement of an isotropic model); such renditions are shown in Supp Figs S4a and S4b for the *P*1 and *P*6 structures, respectively. Analogously, Supp Figs S4c and S4d provide thermal ellipsoid representations of the patterns of variation in anisotropic ADPs across the dodecamer and monomer. In both sets of depictions, Figs S4a/b and S4c/d, colours are graded by the magnitude of per-atom *B*_eq_ values, from low (blue) to medium (white) to high (red). To initially assess the relative contributions of static disorder (e.g., variation in rotameric states across subunits) and dynamic disorder (e.g., harmonic breathing modes and other collective/global motions) in variable regions such as loop L4 and the termini, a normal mode analysis was performed on a coarse-grained representation of the *Aae* Hfq structures, using an anisotropic network model of residue interactions (see Methods). Illustrative results for the dodecamer and monomer are shown in Supp Figs S4e and S4f, respectively. The pattern of normal mode displacements for both the dodecamer and monomer do not implicate loop L4 in any especially high-amplitude, low-frequency modes (Supp Fig S4f), suggesting that L4’s increased ADPs (elevated *B*_eq_ values) stem more from static disorder rather than any particular dynamical process involving this loop region (though anharmonic dynamics remains possible). The dodecamer calculation does reveal a significant harmonic mode corresponding to anti-symmetric rotation of the two Hfq rings with respect to one another (PE 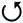, DE 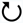; Supp Figs S4e). This result is consistent with our observation that the only large-scale (do-decamer-scale) structural difference between the two rings is a slight rotation of one relative to the other, versus, for instance, a rigid-body tilt (Fig 5, Supp Fig S6a).

At the Hfq ring and supra-ring levels, the refined *P*1 structure reveals an *Aae* Hfq dodecamer consisting of two hexameric rings stacked in a head-to-tail orientation (Fig 5). Propagated across the lattice, this arrangement gives cylindrical tubes with a defined polarity. The tubes run along the crystallographic 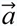 axis, and their lateral packing yields near–six-fold symmetry along this direction; a slight translational shift of the dodecamers in adjacent unit cells, in the plane perpendicular to 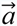, causes the rings to be slightly offset with respect to the lattice tubes (the tubes are not perfectly cylindrical, insofar as the 6-fold axis of an individual Hfq ring is not coaxial with the principal axis of its parent tube). In the dodecamer, the distal face of one Hfq ring is exposed (termed the *DE ring*), while the other ring features a proximal-exposed face (the *PE ring*, Fig 5, right). The *N*-termini of the *DE* hexamer contact the L2-loop/β2-strand region of the *PE* ring, as illustrated in Fig 5 (the L2 loops mark the beginning of strand β2; see the label in Fig 6a). As apparent in the axial view of Fig 5 (left), one ring is slightly rotated relative to the other. Geometric analysis of this rotation (denoted ‘Δ’ in Fig 6a), as well as other rigid-body transformations relating the two rings (Supp Fig S3a, b), shows that the 6-fold symmetry axes of the rings in the dodecamer are not perfectly parallel—a slight tilt occurs between the rings (‘δ’ in Fig 6a). This tilt appears to stem largely from structural differences in the *N*-terminal regions (Supp Fig S3). Consistent with this observation, the set of six *N*-terminal regions of the *DE* ring (which mediate ring⋯ring interactions within a dodecamer) exhibit higher *B*_eq_ values and greater conformational variability than do the *N*-termini of the *PE* ring (which mediate do-decamer⋯dodecamer contacts between unit cells), as can be seen in Supp Fig S4a.

Noncovalent molecular interactions between the proximal⋯distal faces mediate the association of Hfq rings into a dodecamer, and a slightly altered (translationally shifted) version of these same energetically favourable interactions stiches together the dodecamers into a set of crystal lattice contacts in the *P*1 form of *Aae* Hfq. Notably, a *proximal*→*distal* stacking geometry is the chief mode of ring association in the *Aae P*6 lattice too. *Aae* Hfq dodecamers clearly occur in the *P*1 lattice, with a substantial amount of buried surface area (BSA) defining the PE•DE ring interface (Fig 5). Specifically, 3663 ± 244 Å^2^ of SASA is occluded between the *PE* and *DE* hexamers. Note that this quantity is being reported as *BSA* = *ASA*_*PE*_ + *ASA*_*DE*_ − *ASA*_*PE*•*DE*_, where *ASA*_*i*_ is the ASA of species *i*, rather than as the per-subunit value (which would be given by half of the above expression, and which would assume perfect 2-fold symmetry of the interface); also, note that this *mean* ± *standard deviation* is reported from the results of five different surface area approaches, as mentioned in the Methods.

### 3.4. Crystal structure of *Aae* Hfq bound to U_6_ RNA

Upon co-crystallization with U_6_ RNA a second, distinct *Aae* Hfq crystal form could be indexed in *P*6, with unit cell dimensions of *a* = *b* = 66.19 Å, *c* = 34.21 Å. In this form, the cell geometry, solvent content and molecular mass of *Aae* Hfq are only compatible with a single Hfq monomer/ASU, and the crystallographic 6-fold was presumed to generate intact hexamers such as shown in Fig 7a. Specifically, co-crystallization of *Aae* Hfq with this model uridine-rich RNA was achieved by incubating purified Hfq samples with 500 µM U_6_ RNA prior to crystallization trials. The complex crystallized in 0.1 M sodium cacodylate, 5% w/v PEG 8000 and 40% v/v MPD, and the denaturant compound Gnd was found to be an effective additive (Supp Table S2). The crystal structure of the *Aae* Hfq•U_6_ RNA complex was refined to 1.50 Å resolution (Fig 7); we emphasize that the initial solution of this structure was achieved independently from the *apo P*1 form, via molecular replacement, using *P. aeruginosa* Hfq as a search model.

**Figure 7.**
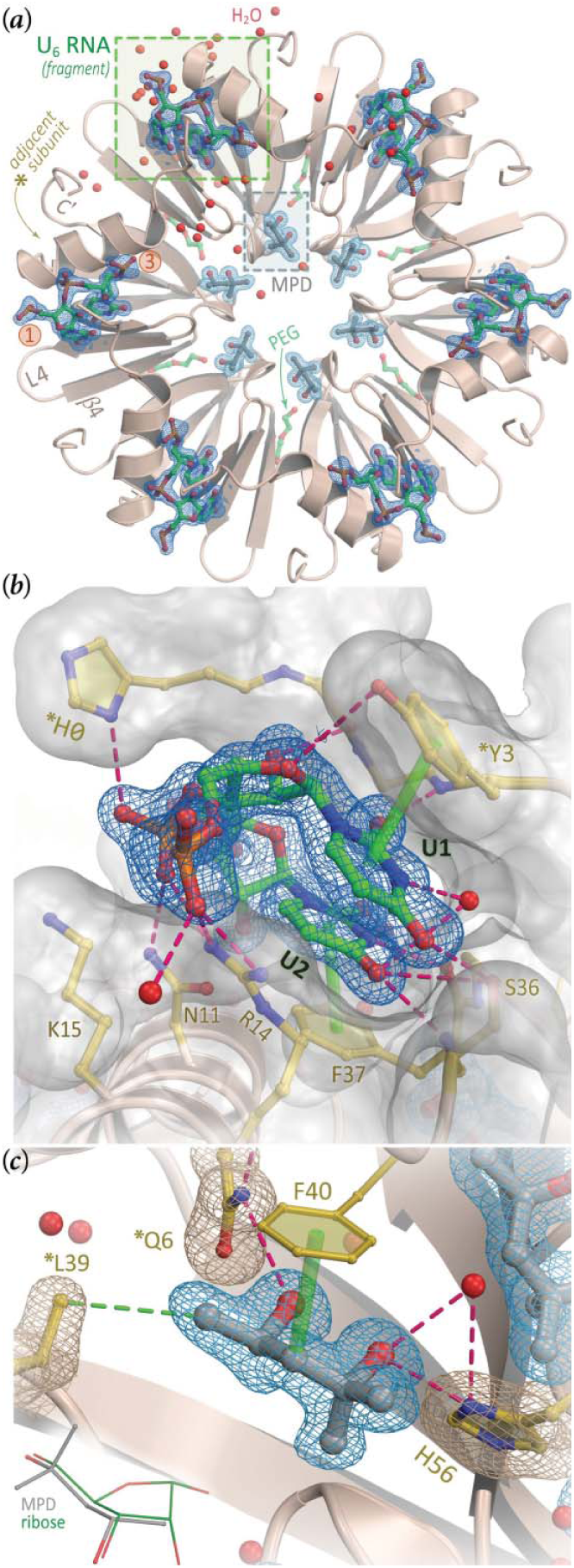
Crystal structure of *Aae* Hfq with U–rich RNA bound at the lateral rim. The asymmetric unit of the *P*6 form contains a single Hfq subunit, shown as a tan-coloured ribbon diagram (*a*), in addition to 36 H_2_O molecules (red spheres), a molecule of PEG (lime-coloured carbons), a molecule of MPD (gray carbons) and one molecule of U_6_ RNA (green carbons). Non–protein atoms are represented as balls-and-sticks, using CPK colours (except as noted above for carbons). Expansion of the ASU to the full *P*6 cell gives an intact Hfq hexamer, shown onto the proximal face in (*a*). The meshes delimit the 2*mF*_*o*_ − *DF*_*c*_ electron density map, contoured at 1.5*σ* and shown only in the regions of RNA (dark blue) or MPD (light blue). The fragment of U_6_ that could be unambiguously built into electron density contained two complete uridines and the 5′ phosphate moiety of the next residue; the path of this RNA strand is denoted by orange ① and ③ symbols for the ribonucleotides, from 5′→3′. Unexpectedly, U_6_ nucleotides were found on the outer rim of *Aae* Hfq, in a position analogous to the lateral site of other Hfqs (*b*), while a molecule of MPD occupied the U-rich–binding pore as shown in (*c*). This magnified view (*b*) of the lateral site (same colour scheme as *a*) shows the RNA-contacting residues (labelled) in greater detail; asterisks distinguish residues from the *N*-termini of a neighbouring subunit, as also indicated in (*a*). Electron density maps such as this one were readily interpretable as RNA (see also Supp Fig S5). The magenta dashed lines (hydrogen bonds) and semi-transparent green cylinders (π-stacking interactions) indicate enthalpically favourable Hfq⋯RNA contacts. Most such contacts are mediated by both backbone and side-chain atoms of *Aae* Hfq, as well as the nucleobase and phosphodiester groups of the RNA; the ribose rings project outward from the cleft, and interact with Hfq more sparsely. (*c*) MPD binds at the pore and mimics the Hfq⋯uridine contacts found at the proximal RNA-binding site in some Hfq homologs. Contacts denoted by magenta dashed lines identically match the contacts to a uridine nucleotide in other Hfq structures containing U-rich RNA (see also Supp Fig S6). The green line indicates a van der Waals contact between L39 and MPD, and the green cylinder denotes another apolar interaction between *Aae* Hfq⋯MPD; this latter contact would presumably be replaced by a π-stacking interaction between F40 and a uracil base, for a U-rich RNA putatively bound at the proximal site.

The residues that are crucial in forming the proximal (U-rich) RNA–binding pocket in *E. coli* and other Hfq homologs (Q8, F42, K56, H57) are conserved in the *Aae* Hfq sequence (Fig 1), which led us to anticipate that any bound U_6_ would be localized to the proximal pore region. Instead, a molecule of the MPD cryo-protectant was found to occupy the proximal site of the Hfq hexamer, with the MPD hydroxyl groups hydrogen-bonded to the sidechains of *Aae*’s H56 and *Q6 residues (Fig 7c); in addition, the MPD makes van der Waals contact with other conserved residues that line the proximal site (*L39, F40). During refinement of this structure, two nucleotides of the U_6_ RNA molecule, including the flanking 5′ and 3′ phosphates (the latter coming from the third U), were readily discernible in *mF*_*o*_ − *DF*_*c*_ difference electron density maps (Fig S5). Notably, processing and reduction of the diffraction data from the *P*6 form in *P*1 yielded similar electron density for the RNA at each lateral binding pocket in the hexamer (Fig S5). Rather than being bound at the proximal site, the uridine residues of U_6_ were found in a cleft formed between the *N*-terminal α-helix and strand β2, in a position located roughly near the outer rim of the *Aae* Hfq toroid (Fig 7a).

### 3.5. RNA binding at the outer rim of the *Aae* Hfq hexamer: Structural details

The *Aae* Hfq•U_6_ structure reveals a lateral binding pocket that accommodates two nucleotides of uridine. The *N*-terminal α-helix primarily contacts the phosphodiester and ribose groups, and the β2 strand interacts mostly with the uracil bases (Figs 7a, 7b, 8a). As a consequence of this RNA-binding geometry, both nucleotides that were fully built into electron density (U1, U2) are held in a bridging, *anti*-conformation (χ = −165.2° for U1, −116.8° for U2), with the ribose moieties extending outward from the pocket (Fig 7b). Interestingly, while the U1 ribose is in the 3′-*endo* conformation typically seen in canonical (*A*-form) RNA structures, with a pseudo-rotation phase angle (*P*) of 17.5º for this South pucker, the U2 sugar adopts a less typical 2′-*endo* conformation (*P* = 163.2º).

**Figure 8.**
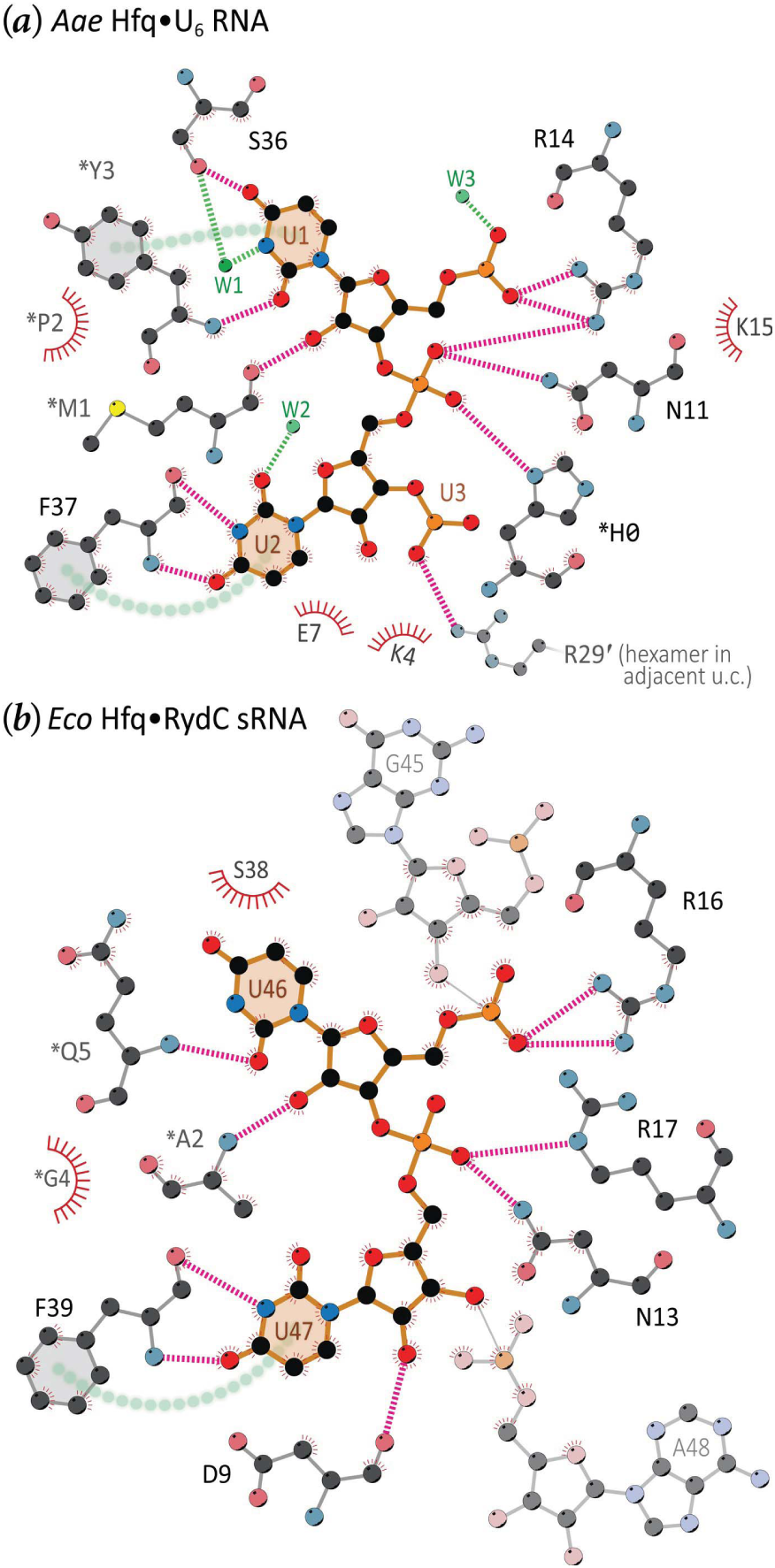
Conserved pattern of interatomic contacts at the lateral RNA-binding site of Hfq hexamers. In this schematic diagram of the interatomic contacts between the lateral site of *Aae* Hfq, U_6_ RNA and nearby H_2_O molecules (*a*), protein atoms are shown as ball-stick representations (CPK colouring, light grey carbons) and covalent bonds in the nucleotides are drawn as thicker, orange-coloured lines. For clarity, only a subset of H_2_O molecules is drawn (green, labelled ‘W#’). Here, asterisks denote another Hfq chain in the same unit cell and the prime symbol denotes a neighbouring cell. Hydrogen bonds are magenta for protein⋯RNA interactions, while those to H_2_O are green. Stacking interactions between aromatic entities φ_1_ and φ_2_ are indicated by green circles from φ_1_⋯φ_2_. Two nucleotides of uridine (labelled) appear in an open, bridging conformation with the α-helix and β2 strand of an Hfq monomer (grey flanking regions). The phosphate groups are hydrogen bonded to N11 and R14 of the *N*-terminal α-helix, while the nucleobase hydrogen bonds with the backbone atoms of strand β2 (specifically, S36 and F37), thus imparting specificity for uridine. Note that additional π-stacking interactions are present between the side-chain of F37 and RNA base U2, as well as within the RNA (between U2⋯U1). (*b*) The lateral pocket of *Eco* Hfq is shown, complexed with the sRNA RydC (same colouring scheme and conventions as in *a*). The U46 and U47 bases adopt conformations similar to those seen in *a*, with the phosphate groups contacting residues of the α-helix. F39 π-stacks with U47, analogous to the interaction seen in *Aae* Hfq. Note that the adjacent G45 and A48 bases are flipped away from the pocket, and are shown here to offer context in the overall sequence of the sRNA. While not strictly conserved in terms of precise amino acid sequence, the *N*-terminal regions of the *Aae* and *Eco* Hfq homologues do provide similar backbone interactions with U1 and U46, respectively. Note also the directionality of the RNA backbone, which follows the same 5′→3′ path along the lateral site on the surface of the *Aae* and *Eco* Hfq rings (see also Figs 7a, b).

Protein⋯RNA interactions are mediated by both side-chain and backbone atoms of *Aae* Hfq. The full set of interactions is shown in 3D in Fig 7a, b, and schematically in Fig 8a. Two side-chains in *Aae*’s *N*-terminal α-helix, N11 and R14, contact the phosphodiester groups (denoted ‘Ⓟ’ for brevity), and another cationic residue (K15) is 3.6 Å from the Ⓟ between the two uridines. Backbone and side-chains atoms from strand β2 hydrogen-bond with the bases, ensuring uridine specificity (Figs 7b, 8, 9). In particular, both the carbonyl oxygen and amide nitrogen of F37 interact with N3 and O4 of U2, respectively, while the hydroxyl side-chain of S36 contacts the exocyclic O4 of U1. S36 also helps position a pivotal H_2_O that directly hydrogen-bonds to both the N3 atom of U1 and the S36 hydroxyl (Fig 8a); this well-ordered (ice-like) water molecule engages in a network of hydrogen-bonds, in a distorted tetrahedral geometry (additional structural waters also contact the uracil and Ⓟ moieties, as shown in Fig 8). Other interactions at the lateral site include a series of three π-stacking interactions (Fig 8a): between the phenyl ring of F37⋯U2, the U1⋯U2 bases, and the phenolic ring of *Y3⋯U1. RNA-binding at the lateral site is composite in nature, involving not just residues of strand β2 and helix α1 of one subunit, but also the *N*-terminal tail of an adjacent subunit in the ring. The irregularly structured *N*-terminal tail of one Hfq monomer extends into the neighbouring lateral site, where the *N*-terminal sequence H^0^M^1^P^2^Y^3^K^4^ nearly ‘covers’ that rim site and supplies additional contacts with RNA. For instance, *Y3 engages in the π-stacking mentioned above, as well as a hydrogen-bond between its amide nitrogen and the O2 of U1 (an interaction that does not select between uracil and cytidine). Also in this region, the backbone carbonyl oxygen of *M1 hydrogen-bonds to the ribose O2′ of U1, thus contributing to discrimination between RNA and DNA. Finally, we note that two contacts in this region may be spurious: (*i*) the ^*^H0⋯Ⓟ interaction, where residue ^*^H0 is from the recombinant construct (not wild-type *Aae* Hfq; see numbering in Supp Fig S1); and (*ii*) the R29′⋯Ⓟ interaction, which is a crystal lattice contact (the prime symbol on R29′ indicates an adjacent unit cell).

**Figure 9.**
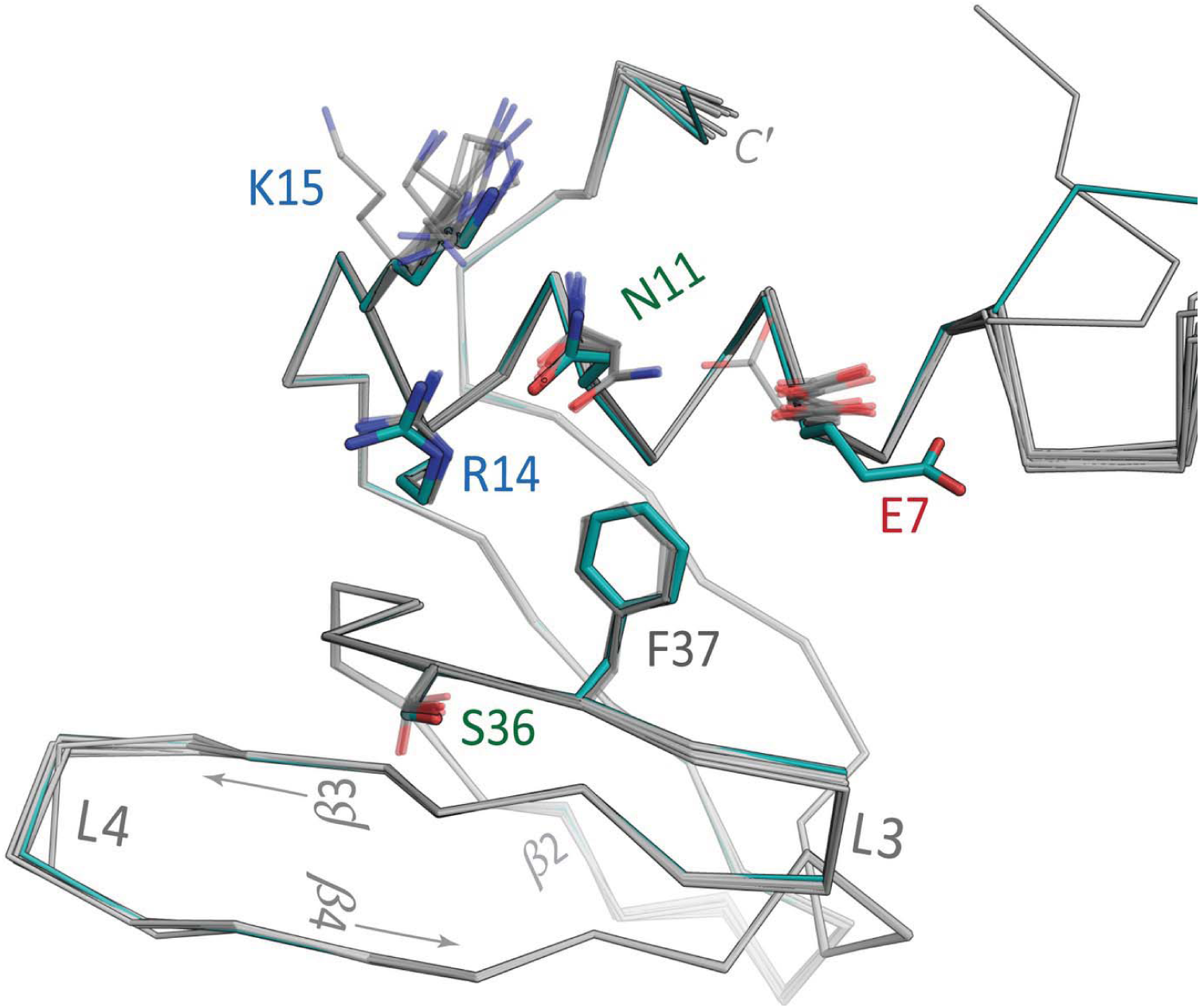
The lateral site of *Aae* Hfq is pre-structured for RNA-binding. The 3D structure of the single, unique monomer from the Hfq-U_6_ co-crystal structure (teal backbone) was superimposed with the twelve subunits of the *apo* Hfq structure (grey). Residues that contact RNA, to within ≈ 3.6 Å in the *P*6 Hfq•U_6_ structure, are shown as sticks for both the *P*6 and *P*1 structures. Apart from residue E7, which sterically occludes the binding pocket and thus likely adopts a different conformation upon RNA binding, note that the side-chains in the *apo* structure adopt rotameric states quite similar to those in the 3D structure of U_6_-bound *Aae* Hfq. This finding suggests pre-organization of *Aae* Hfq’s RNA-binding site.

Comparison of the *Aae* Hfq•U_6_ structure with the independently-refined *apo Aae* Hfq structure suggests that the lateral RNA-binding site is essentially pre-structured for RNA complexation (Fig 9). In terms of comparative structural analysis, note that the *apo/P*1 and RNA-bound*/P*6 structures are at equally high resolutions (1.49, 1.50 Å respectively; Table 1), were refined in similar manners (e.g., using anisotropic ADPs), and are of comparable quality in terms of *R*_work_/*R*_free_, stereochemical descriptors, etc. (Table 2). Residues N11, R14, S36 and F37, which are phylogenetically conserved to varying degrees (Fig 1), largely define the structural and chemical topography of the lateral site (Fig 7a). As shown in Fig 9, these crucial residues adopt nearly identical rotameric states in the *apo* and U_6_–bound forms of *Aae* Hfq. The two principal RNA-related structural differences, in going from the *apo* to the U_6_–bound forms, are: (*i*) a shift in the residue E7 rotamer (Fig 9, red label), positioning this side-chain away from the pocket and thus enabling the U2 base to be accommodated, and (*ii*) the precise path of the *N*-terminal tail (i.e., the ≈5 residues preceding helix α1), which varies with respect to the lateral site. In the dodecameric *apo* structure, six of the *N*-termini mediate ring⋯ring contacts (Fig 5) while the other half mediate lattice contacts, giving rise to one source of structural heterogeneity in this region. In terms of intrinsic flexibility, normal mode calculations (Supp Fig S4 and Methods) indicate that the *N*-terminal regions in the hexamer are highly flexible when free in solution, but rigidified (as much as any other part of the Sm fold) when sandwiched between the Hfq rings.

## 4. Discussion

The *apo* form of *Aae* Hfq crystals, refined to 1.49 Å in space-group *P*1, reveals a dodecamer comprised of two hexamers in a head-to-tail orientation. The individual subunits of *Aae* Hfq are similar in structure, with a mean pairwise RMSD less than ≈ 0.3 Å for all monomer backbone atoms. The largest differences among the 13 independently-refined Hfq monomer structures (12 in *P*1, one in *P*6) occur in the *N*-terminal and L4 loop regions; notably, these are the two regions that mediate much of the interface between rings (*distal*⋯*proximal* face contacts in Fig 5), as well as the intermolecular contacts between dodecamers across the lattice. The patterns of structural differences are also captured in the symmetric matrix of pairwise RMSDs between chains: hierarchical clustering on this distance matrix results in the monomers that comprise the *PE* (chains A–F) and *DE* (chains G–L) hexameric rings partitioning into two distinct groups (Fig 6c, Supp Fig S3c).

Sm proteins, including Hfq, exhibit a strong propensity to self-assemble into cyclic oligomers that then crystallize as either (*i*) cylindrical tubes with a defined polarity, via head→tail stacking of rings (*Aae* Hfq and *Mth* SmAP1 are two examples), or (*ii*) head↔head oligomers, often with dihedral point-group symmetry (*Pae* SmAP1 is an example (Mura, Kozhukhovsky*, et al.*, 2003)). An examination of the lattice packing of all known Hfq structures (data not shown) reveals at least one example of each possible ring-stacking mode for a dodecameric assembly: (*i*) a *proximal*•*proximal* interface, as seen in the extensive interface between hexamers of an Hfq orthologue from the cyanobacterium *Synechocystis* sp. PCC6803 (PDB ID 3HFO; (Bøggild *et al.*, 2009)); (*ii*) a *distal*•*distal* interface, observed in *S. aureus* Hfq (PDB ID 1KQ2; (Schumacher *et al.*, 2002)) and in *P. aeruginosa* Hfq, with a more modest interface and relative translational shift of one ring (PDB ID 4MMK; (Murina *et al.*, 2014)); and (*iii*) the *head*→*tail* packing of two rings in a *L. monocytogenes* (*Lmo*) Hfq structure, in *apo* and RNA-bound forms (PDB ID 4NL2; (Kovach *et al.*, 2014)). The *Aae head*→*tail* interface (Fig 5) is tighter than that between the *Lmo* Hfq rings, but otherwise the stacking in these two Hfq structures resemble one another even in fine geometric details (e.g., the top/bottom, *PE/DE*, rings are similarly rotated with respect to one another). Also, the *S. aureus distal*•*distal* dodecamer buries 2666 Å^2^ of surface area, which is considerably less than the ≈ 3700 Å^2^ of ΔSASA determined here for *Aae* Hfq’s *distal*•*proximal* stacking mode.

As a point of reference, note that the above ΔSASA quantities represent less buried surface area than in the *ring•ring* interfaces found in the structures of various Sm and SmAP homologs. (Recall that Hfq rings are hexameric while SmAPs are generally heptameric, meaning that there likely will be a systematic difference in ΔSASA trends simply by virtue of subunit stoichiometry.) The *ring•ring* interfaces in *P. aerophilum* and *M. thermautotrophicum* 14-mers occlude 7550 and 3000 Å^2^, respectively. Unlike *P. aerophilum* SmAP3, where the burial of >21,000 Å^2^ along an intricate interface between stacked rings suggests *bona fide* higher-order oligomers (Mura, Phillips*, et al.*, 2003), the extent of the *Aae* Hfq *distal•proximal* interface does not as clearly indicate whether or not dodecamers exist. The free energy of association between *Aae* Hfq’s *PE•DE* rings, 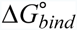, can be estimated via the linear relationship 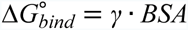 (the slope, *γ*, is often taken as ≈ 20-30 cal∙mol^-1^∙Å^-2^ (Janin *et al.*, 2008)); however, *Aae* Hfq’s *PE*•*DE* interface is not primarily apolar in character, so this approach may severely overestimate the 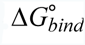. Also, in terms of the existence and potential relevance of higher-order states, recall that *Aae* Hfq can form dodecamers *in vitro*, at least when bound to an A– rich RNA and assayed by AnSEC (Fig 3b, blue arrow). Nevertheless, despite all these observations, *(i)* whether or not Hfq dodecamers actually occur *in vivo*, beyond crystalline and *in vitro* milieus (such as in AnSEC experiments) remains unclear, and (*ii*) even if such dodecamers do exist, the potential physiological activities and functional roles of higher-order oligomeric states of Hfq remains murky.

Intriguingly, our solution-state AnSEC data are consistent with the binding of A_18_, presumably at the distal face of (Hfq)_6_, causing a shift in the distribution of *Aae* Hfq oligomeric states from hexamers (only) to a more dodecameric population (Fig 3). This effect may be attributed to the longer A_18_ strand simultaneously binding to two Hfq rings, giving a ‘bridged’ ternary complex. There also appears to be some length-dependence to the interaction of A-rich RNAs with Hfq, as we found that A_6_ did not exhibit high-affinity binding to *Aae* Hfq; this dependence may stem from mechanistic differences in the early (initiation) stages of the kinetic mechanism for Hfq⋯RNA binding. *Aae* Hfq demonstrates a nanomolar affinity for A_18_ and U_6_ RNA that is selective (C_6_ does not bind) and that is consistent with the properties of Hfq homologs characterized from other bacteria, both Gram-negative (e.g., proteobacteria such as *E. coli*) and Gram-positive. For instance, the magnesium–dependence of the *Aae* Hfq•U_6_ interaction (Fig 4), with 10-fold stronger binding in the presence of Mg^2+^, mirrors the Mg^2+^-dependency of U-rich–binding by Hfq homologs from the pathogenic, Gram-positive bacterium *Listeria monocytogenes* (*Lmo*) and the Gram-negative *E. coli* (*Eco*; (Kovach *et al.*, 2014)). For both *Lmo* and *Eco* Hfq, the inclusion of 10 mM magnesium increased the U_6_-binding affinity by >100-fold; the effect was similar, but less pronounced, for U_16_ (≈ 3-4–fold increase). Thus, the Mg^2+^-dependency of the *Aae* Hfq•U_6_ RNA interaction is intermediate between these two extremes.

At present, only two other known Hfq structures contain a nucleic acid bound to the lateral site. These structures are: (*i) Pae* Hfq co-crystallized with the nucleotide uridine–5′–triphosphate (UTP; PDB ID 4JTX; (Murina *et al.*, 2013)) and (*ii) Eco* Hfq bound to a full-length sRNA known as RydC (PDB ID 4V2S; (Dimastrogiovanni *et al.*, 2014)). Comparison of the lateral RNA-binding sites of the *Aae*, *Pae* and *Eco* Hfq structures reveals a highly conserved pocket formed by N13, R16, R17, S38 and F39 (*Eco* numbering; see also Fig 1). In *Aae* Hfq, K15 appears to be homologous to *Eco* R16, insofar as this side-chain is well-positioned to engage in electrostatic and hydrogen bond interactions with an the sugar-phosphate backbone of a bound RNA (Figs 7b, 8, 9). This structural feature can be seen both in *Eco* Hfq (R17 with the phosphate of a neighbouring nucleotide) and in *Pae* Hfq (K17 with the 5′ phosphate tail of UTP). Notably, uridine is the only nucleotide that has been found to bind at the lateral site in all three of these Hfq structures—*Eco*, *Pae*, and now *Aae*.

At a resolution of 1.5 Å, the *Aae* Hfq•U_6_ structure offers new insights into the apparent specificity of the lateral pocket for uridine nucleosides. We see that interactions with the backbone of strand β2 provide discrimination between uridine and cytidine bases in the cognate RNA. One uracil base πstacks with a key phenylalanine residue, while the second uracil stacks atop the preceding nucleobase. The second nucleotide adopts a C2′-*endo* conformation, leading to accommodation of the base in the binding pocket on Hfq’s surface. In this configuration, the *N*-terminal region may then provide further enthalpically favorable interactions that stabilize the complex. The *Aae* Hfq lateral site includes two of the three arginine residues of the ‘arginine patch’, known to be important for annealing of sRNAs and mRNAs (Panja *et al.*, 2013). We propose that the third arginine of this motif acts primarily electrostatically—without directionality, and non-specifically as regards RNA sequence—in order to enhance the diffusional association of an RNA by ‘guiding’ it towards the lateral pocket. In addition, the physicochemical basis for the phylogenetic conservation of the lateral site may be that it simply provides additional surface area for Hfq⋯sRNA interactions, perhaps supplying an extended platform for the ‘cycling’ of RNAs across the surface of the Hfq ring (Wagner, 2013); similarly, the rim site may serve as an additional ‘anchor’ for the association of moderate-length, U-rich RNAs that bind with low intrinsic affinity for the proximal site, but which can reach the lateral/rim site. We propose that the lateral site, which is structurally well-defined on the outer rim of the *Aae* Hfq hexamer, is a biologically relevant region that functions in binding (U)_*n*_ segments of RNA containing at least two consecutive uridine nucleotides; moreover, we propose that this RNA-binding region is conserved in even the most ancient bacterial lineages.

The structural features of Hfq⋯RNA interactions in homologs from evolutionarily ancient bacteria share some similarity with the properties of Sm-like archaeal proteins (SmAPs), such as a SmAP from the hyperthermophile *Pyrococcus abyssi* (*Pab*) that was co-crystallized with U_7_ RNA (Thore *et al.*, 2003). Interestingly, the oligoribonucleotide in that crystal structure was found in two sites: the canonical U-rich–binding site near the lumen of the ring (analogous to Hfq’s proximal site), as well as a ‘secondary’ pocket on the same (proximal) face. This secondary site of *Pab* SmAP is distant from the U–binding site, lying between the *N*-terminal α-helix and strand β2 of the Sm fold. Note that the ‘lateral site’ of Hfq had not yet been discovered as an RNA interaction region at the time of the *Pab* structure determination. The secondary RNA-binding site in *Pab* SmAP also contains a phenylalanine residue that is conserved among Hfq homologs and that is required for π-stacking with the nucleobase. However, the asparagine residue found at the lateral site of all characterized Hfq homologs is instead a histidine in *Pab* SmAP; this residue’s imidazole side-chain provides an additional stacking platform for an adjacent ribonucleotide in the *Pab* complex, in an interaction that is unseen with known Hfq homologs. The α-helix of *Pab* SmAP does not extend as far as that of Hfq, and the arginine-rich patch that occurs at this rim area in Hfq homologs is but a single lysine residue in *Pab* SmAP. Nevertheless, the presence of this partially conserved lateral pocket in *Pab* SmAP does suggest an ancient, common origin for this mode of protein⋯RNA recognition by Hfq and other members of the Sm superfamily. Somewhat similarly, a uridine-binding site was crystallographically identified in *Pyrobaculum aerophilum* SmAP1, in a region on the ‘L3 face’ (analogous to Hfq’s proximal face) that lies distal to the canonical U–rich RNA-binding site at the inner surface of the pore; this L3-face region was described as a ‘secondary’ binding site because of relatively weak electron density (Mura, Kozhukhovsky*, et al.*, 2003). We can now see that the secondary U-rich–binding sites in at least two archaeal Sm proteins, from *Pab* and *P. aerophilum*, occupy a region that is roughly analogous to the lateral rim of Hfq.

The historical lack of structural data on RNA-binding at the Hfq lateral site may be because uridine-rich RNAs—such as might localize to the lateral rim—are also capable of binding to the higher-affinity proximal site. A single binding event is consistent with the idealized shape of our *Aae* Hfq•U_6_ binding curves (Fig 4a), which bear no hint of multiple transitions or non-two-state binding. This could indicate that U_6_ binding at the proximal and distal sites differs by at least an order of magnitude (beyond the detection range of our assay). For the *Aae* Hfq•U_6_ complex reported here, we suspect that two facets of our crystallization efforts serendipitously shifted the RNA–binding propensity towards the lateral site. First, MPD was present at high concentrations in our crystallization condition (many Hfq homologs reported in the literature were crystallized with PEGs, not MPD). MPD is a commonly-used precipitating agent and cryoprotectant, and inspection of electron density maps reveals it to be associated with all 12 subunits of the *apo* form of *Aae* Hfq; specifically, 24 of the 25 MPDs found in the *P*1 electron density are bound in one of two locations (Fig 5), one of these being the proximal RNA-binding site. Moreover, an MPD molecule was also bound in the *P*6 (U_6_-bound) crystal forms, in clear density at the proximal site (Fig 7). In terms of structural and chemical properties, the hydroxyl groups of MPD closely mimic the ribose and uracil moieties of uridine, as shown in Supp Fig S6. Residues H56 and Q8 have been identified as two key residues in the proximal site that contact the ribose 2′-OH and uracil’s exocyclic O2 atom upon binding of U_6_ at the proximal site (Schumacher *et al.*, 2002). However, in our *Aae* Hfq structure these two residues instead contact MPD (*Q6 and H56 in Fig 7c). The lateral RNA-binding site, however, does not include many contacts to ribose (versus the phosphate and nucleobase groups), and thus MPD would not be expected to compete as strongly against RNA-binding at that site. The second unique feature of *Aae* Hfq that may increase affinity for U-rich RNA at the lateral site is the flexible *N*-terminal tail, which folds over the lateral site when nucleic acid is bound, further stabilizing the bound U_6_ RNA. In our work, the *N*-terminus includes three plasmid-derived residues that remain after cleavage of a His6× tag used in protein purification (G^−2^S^−1^H^0^; Supp Fig S1a). The additional histidine contacts the phosphate of nucleotide U2 (Figs 7b, 8). In addition, the native sequence includes a tyrosine residue that provides further aromatic stacking interactions with base U1 (residue *Y3 in Figs 7b, 8). This tyrosine residue is not conserved among other Hfq homologs, many of which contain a glutamate at this position (Fig 1).

The crystallographic and biochemical work reported here reveals that the putative Hfq homolog encoded in the *A. aeolicus* genome is an authentic Hfq, as it (*i*) adopts the Sm fold, (*ii*) self-assembles into hexameric rings that can associate into higher-order double rings in the lattice (as do many known Hfqs), and (*iii*) binds A/U-rich RNAs with high affinity (and selectivity). Perhaps most exciting, these structural and functional properties are recapitulated by an Hfq homolog from the Aquificae phylum, which may be the most basal, deeply-branching lineage in the *Bacterial* domain of life (Bocchetta *et al.*, 2000, Burggraf *et al.*, 1992). To date, all Hfq structures have been limited to three phyla: (*i*) most Hfq structures are from the *Proteobacteria*, (*ii*) a few are from the (mostly Gram-positive) *Firmicutes* and, finally, (*iii*) two known homologs are of *Cyanobacterial* origin. Because of its basal phylogenetic position, the *Aae* Hfq structures reported here—the first Hfq structures from outside these three bacterial lineages—suggest that members of the Sm/Hfq superfamily of RNA– associated proteins, along with at least some of their RNA–binding properties, likely existed in the last common ancestor of the *Bacteria*.

## Acknowledgements

We thank H. Huber (Regensburg) for providing a sample of *A. aeolicus* genomic material; J. Bushweller (UVa) for access to a fluorescence plate reader; J. Shannon (UVa) for assistance with MALDI-TOF instrumentation; D. Cascio and M. Sawaya (UCLA) for crystallographic advice; and L. Columbus (UVa) and C. McAnany (UVa) for helpful discussions. Beamlines NE-CAT 24-ID-C/E at Argonne National Lab’s *Advanced Photon Source* are DOE facilities (DE-AC02-06CH11357), with NIH funding for general operations (GM103403) and for the Pilatus detector (RR029205). This work was funded by the Jeffress Memorial Trust (J-971), NIH training grant T32GM080186 (P.S.R.) and NSF CAREER award MCB–1350957.

